# scEGOT: Single-cell trajectory inference framework based on entropic Gaussian mixture optimal transport

**DOI:** 10.1101/2023.09.11.557102

**Authors:** Toshiaki Yachimura, Hanbo Wang, Yusuke Imoto, Momoko Yoshida, Sohei Tasaki, Yoji Kojima, Yukihiro Yabuta, Mitinori Saitou, Yasuaki Hiraoka

**Affiliations:** Mathematical Science Center for Co-creative Society, Tohoku University, Sendai 980-0845, Japan; Department of Anatomy and Cell Biology, Graduate School of Medicine, Kyoto University, Yoshida-Konoe-cho, Sakyo-ku, Kyoto 606-8501, Japan; Institute for the Advanced Study of Human Biology, Kyoto University Institute for Advanced Study, Kyoto University, Yoshida-Konoe-cho, Sakyo-ku, Kyoto 606-8501, Japan; Faculty of Science, Kyoto University, Kitashirakawa Oiwake-cho, Sakyo-ku, Kyoto 606-8502, Japan; Department of Mathematics, Faculty of Science, Hokkaido University, Sapporo 060-0810, Japan; Center for iPS Cell Research and Application, Kyoto University, 53 Kawahara-cho, Shogoin, Sakyo-ku, Kyoto 606-8507, Japan

**Keywords:** trajectory inference, optimal transport, Gaussian mixture model, single-cell biology, epigenetic landscape

## Abstract

Time-series scRNA-seq data have opened a door to elucidate cell differentiation, and in this context, the optimal transport theory has been attracting much attention. However, there remain critical issues in interpretability and computational cost. We present scEGOT, a comprehensive framework for single-cell trajectory inference, as a generative model with high interpretability and low computational cost. Applied to the human primordial germ cell-like cell (PGCLC) induction system, scEGOT identified the PGCLC progenitor population and bifurcation time of segregation. Our analysis shows *TFAP2A* is insufficient for identifying PGCLC progenitors, requiring *NKX1-2*. Additionally, *MESP1* and *GATA6* are also crucial for PGCLC/somatic cell segregation. These findings shed light on the mechanism that segregates PGCLC from somatic lineages. Notably, not limited to scRNA-seq, scEGOT’s versatility can extend to general single-cell data like scATAC-seq, and hence has the potential to revolutionize our understanding of such datasets and, thereby also, developmental biology.

## Background

The “epigenetic landscape” proposed by the renowned biologist C. H. Waddington is a well-known metaphor for describing cell differentiation and is a key concept in developmental biology [1]. In this conceptual model, cells begin as stem cells at the top of this landscape and differentiate into more specialized cell types as they move down the valleys during the development, with the ridges representing potential barriers that prevent transitions between cell types.

Although a useful concept, the actual shapes of the landscapes during differentiation processes have remained unclear in many biological systems. However, recent advances in genome-scale high-dimensional single-cell technologies, such as single-cell RNA sequencing (scRNA-seq) [2, 3], have opened an avenue for inferring the dynamics of cell differentiation in a data-driven manner, as well as for reconstructing Waddington’s landscape. This has made trajectory inference for cell differentiation a central topic in current single-cell and systems biology [4, 5, 6].

Many methods [7, 8] for trajectory inference have been developed using a single snapshot of the scRNA-seq data. In spite of the snapshot data, due to the heterogeneity of the cell population, these methods can identify changes in gene expression levels along the pseudo-time [9, 10, 11, 12, 13]. However, the dynamics of overall cell differentiation are very complex, causing the static trajectory inference described above to have obvious limitations [14]. Recently, time-series scRNA-seq data have been used to overcome this difficulty and to gain more insight into the dynamics of cell differentiation. Nevertheless, the destruction of cells at each measurement severely impedes the identification of cell populations between adjacent time points. Trajectory inference methods based on the optimal transport theory have attracted attention in recent years to deal with this issue.

The optimal transport (OT) is a mathematical theory that provides distances and optimal matchings between probability distributions [15, 16, 17]. It has recently been applied in biology [18], especially in single-cell analysis [19, 20, 21], as well as in several other research fields [22]. Among them, Waddington-OT [23] is a well-known method for inferring the cell lineages by applying a static unbalanced optimal transport to the time-series scRNA-seq data. While it can predict cell lineages from the optimal matching of cell populations, since it does not learn the continuous distributions of cell populations (i.e., a non-generative model), we cannot gain much insight into the intermediate states in the differentiation process. It is also known that such optimal transports between cells do not sufficiently reflect those between the cell distributions [24].

On the other hand, optimal transport methods with generative models based on neural networks have also been reported, such as TrajectoryNet [25], JKONET [26], and PRESCIENT [27]. They can generate data in the intermediate states of the differentiation process. However, the neural networks used there introduce black boxes into these methods, making biological interpretation difficult.

In general, the computational cost of solving optimal transport problems, including the above methods, is very high and can be a bottleneck for trajectory inference. GraphFP [28] is a method that addresses this problem by combining dynamic optimal transports on cell state graphs with the nonlinear Fokker–Planck equation. The key to reducing the computational cost is the formulation using inter-cluster optimal transport. Owing to its construction, this method achieves high biological interpretability and low computational cost. However, since it deals with only the cell lineages of the cell clusters, it cannot infer the continuous state of the differentiation process (e.g., transitions during the merging/separation of cell clusters).

In this paper, we present scEGOT, a novel trajectory inference framework based on entropic Gaussian mixture optimal transport (EGOT). It aims to provide a comprehensive trajectory inference framework to infer the dynamics of cell differentiation from time-series single-cell data. The methodology is based on an inter-cluster optimal transport, where clustering and learning of the distributions are performed on cell populations in the gene expression space using the Gaussian mixture model (GMM), and each Gaussian distribution corresponds to a cell type. The main feature of this method is that it has a clear and rigorous correspondence to a continuous optimal transport. Moreover, its computational cost is significantly low owing to the intercluster optimal transport. Accordingly, we can continuously infer the intermediate states of the cell differentiation process at low computational cost. As a comprehensive framework, scEGOT can construct (i) cell state graphs, (ii) velocity fields of cell differentiation (called cell velocity in this paper), (iii) time interpolations of single-cell data, (iv) space-time continuous videos of cell differentiation with gene expressions, (v) gene regulatory networks (GRNs), and (vi) Waddington’s landscape (Fig. 1).

**Figure 1:**
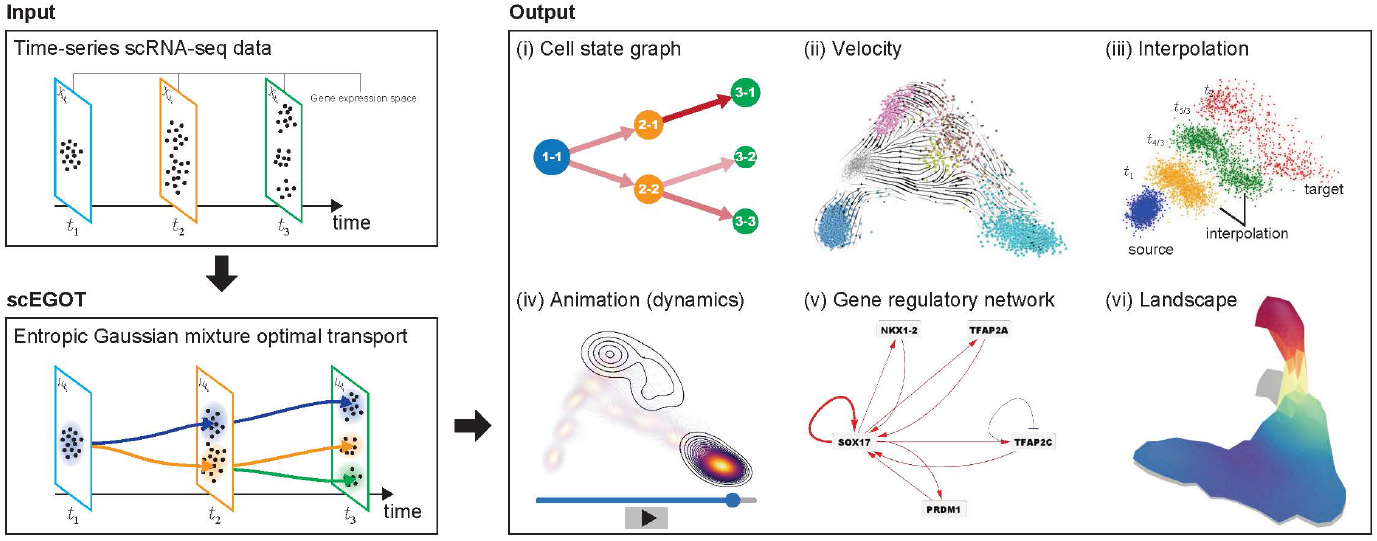
Sketch of the framework of scEGOT. scEGOT is a trajectory inference method based on entropic Gaussian mixture optimal transport (EGOT) that extracts various local and global structures of cell differentiation from scRNA-seq data. scEGOT takes time-series scRNA-seq data as input and outputs the following six cell-differentiation structures: (i) cell state graphs: transitions between cell populations over time; (ii) cell velocity: velocity of cell differentiation in gene expression space; (iii) interpolation: generation of pseudo-scRNA-seq data at intermediate time points; (iv) animation: visualization of gene expression dynamics; (v) gene regulatory network (GRN): regulatory relationships between genes during transitions; and (vi) Waddington’s landscape: cell potency and a global view of the cell differentiation pathway.

As a biological application, we apply scEGOT to the time-series scRNA-seq data of the human primordial germ cell-like cell (human PGCLC) induction system. We demonstrate that scEGOT provides insights into the molecular mechanism of PGCLC/somatic cell segregation. In particular, using the functions of scEGOT, we elucidate the dynamics of the PGCLC differentiation and identify the PGCLC progenitor cell population. Furthermore, we find key genes such as *NKX1-2, MESP1*, and *GATA6* that may play crucial roles during human PGCLC differentiation.

## Results

### Theory of EGOT

We present here the mathematical foundation of EGOT and its application to single-cell biology, called scEGOT. By generalizing [29], the EGOT is formulated by an entropic regularization of the discrete optimal transport, which is a coarse-grained model derived by taking each Gaussian distribution as a single point.

We first summarize the properties of the solution of the EGOT and then discuss the entropic transport plan constructed from the solution of EGOT. Furthermore, we show a correspondence between EGOT and the continuous optimal transport presented by [30]. This theoretical compatibility enables us to present a novel trajectory inference framework in scEGOT. Specifically, this framework allows us to construct a cell state graph and infer the intermediate states and velocity of the cell differentiation process (cell velocity), from which we can further infer the GRNs between adjacent time points and reconstruct Waddington’s landscape in the gene expression space, with high biological interpretability and low computational cost (we refer to *Methods* section for more details).

All proofs of the mathematical statements made here are provided in the supporting information.

### Gaussian mixture model

Let 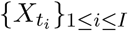 denote the time-series data of the point clouds in ℝ^*n*^(*n* ≥ 1) with *I* time stages (possibly with different sample sizes). For each point cloud 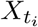, we apply a Gaussian mixture model (GMM) [31]. Then, we obtain

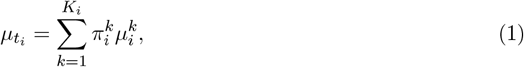

where *K*_*i*_ denotes the number of clusters in the distribution at *t*_*i*_, 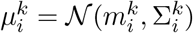 is a Gaussian distribution with a mean vector 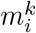 and a covariance matrix 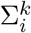, and 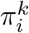 is a non-negative weight of each Gaussian distribution satisfying 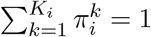. In the scRNA-seq data, 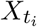 denotes a cell population in the gene expression space ℝ^*n*^ at the time *t*_*i*_, where each cell 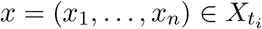 is characterized by the *n* values of the gene expressions, or those in a principal component analysis (PCA) space after the dimension reduction. Then, each Gaussian distribution and its weight are regarded as a certain cell type and its existence probability, respectively.

### Entropic Gaussian mixture Optimal Transport (EGOT)

EGOT connects these Gaussian mixture distributions between adjacent times by solving the following optimization problem for the weights 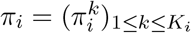 and 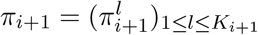

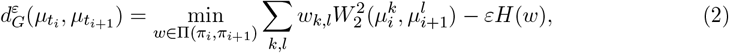

where

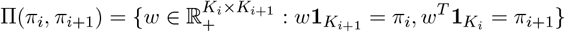

is a subset of *K*_*i*_ ×*K*_*i*+1_ matrices of non-negative real numbers; 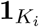 expresses the *K*_*i*_-dimensional vector whose entries are all 1; and (•)^*T*^ denotes the transpose. Here, *W*_2_ is the *L*^2^-Bures– Wassersterin distance [32, 33, 34]

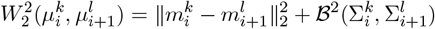

using the squared Bures metric [35]

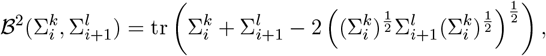

and *H* is an entropy function defined as

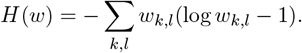

The solution *w* of the EGOT represents how close the Gaussian distributions of the adjacent time points are to each other. Thus, it indicates the similarity between the cell populations when applied to the scRNA-seq data. Since the discrete entropy *H* is a strongly concave function, the objective function of (2) is a strongly convex function. Therefore, the optimization problem (2) has a unique optimal solution. In addition, by considering the Lagrangian associated with the problem (2) and the first-order optimality condition, the following proposition holds:

#### Proposition 1.

*The optimization problem* (2) *has a unique solution w*^*ε*^ *given by*

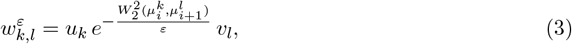

*where* 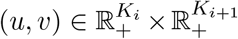 *are derived from the dual variables of the Lagrangian associated with the problem* (2).

We emphasize that by coarse-graining point clouds with Gaussian mixture distributions, the EGOT is significantly less computationally expensive than directly analyzing the optimal transports with full point clouds (e.g., Waddington-OT [23]). Furthermore, as we will see subsequently, EGOT can recover the solution and distance of a continuous optimal transport between Gaussian mixtures, which will provide deeper insights into the stochastic dynamics of cells in the gene expression space.

### Convergence of EGOT and its connection to continuous OT

In this section, we clarify the connection between EGOT and the following continuous optimal transport proposed by [30]:

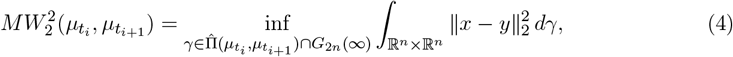

where 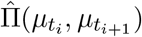 is a set of probability measures on ℝ^*n*^ × ℝ^*n*^ with the given marginals 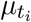 and 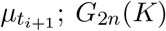 is a set of probability measures on ℝ^*n*^ × ℝ^*n*^, which can be written as Gaussian mixtures with *K* or less components; and 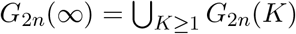.

Let us first consider the convergence of the EGOT (2) and the solution (3) as *ε* → 0. By using an estimate similar to that for discrete entropic regularized optimal transports with Shannon entropy by Weed [36], we can prove the following proposition.

#### Proposition 2.

*The solution w*^*ε*^ (3) *exponentially converges to an optimal transport plan of* 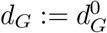 *such that with the maximal entropy as ε* → 0. *That is, there exists M >* 0 *independent of ε >* 0 such that

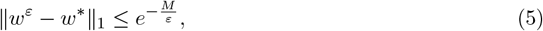

*where w*^*^ *is the solution of*

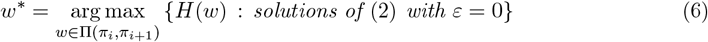

*and* ∥ · ∥_1_ *denotes the* 𝓁_1_ *norm of vectors (viewing w as a vector). Moreover*, 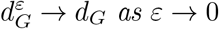 *with the following estimate:*

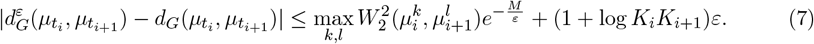

As a result of (5), the solution remains largely unaffected even when the regularization parameter *ε* is changed, and there is almost no impact on the solution.

Next, we define an entropic transport plan

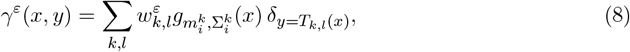

where 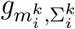 is the multivariate Gaussian distribution

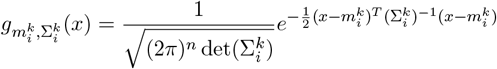

and *T*_*k,l*_ is the optimal transport map between 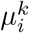 and 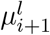 given by

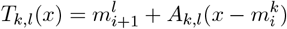

with

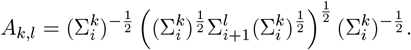

By applying the convergence result (5) and the estimate (7), we obtain the main theorem in this paper, which provides a correspondence between EGOT and the continuous optimal transport (4) (Fig. 2).

**Figure 2:**
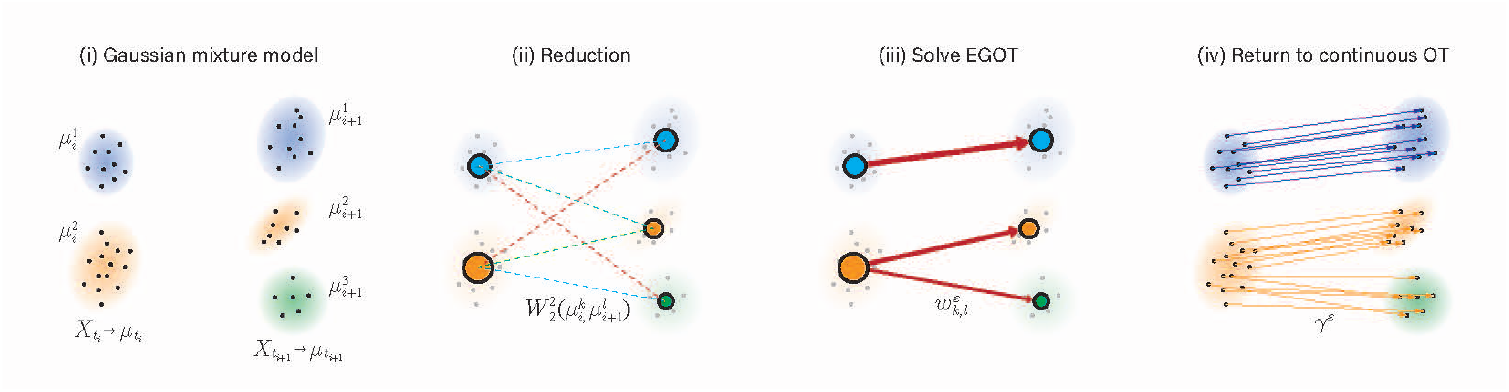
Illustration of the connection between EGOT (2) and continuous optimal transport (4).

#### Theorem 3.

*The continuous optimal transport* (4) *has a minimizer*

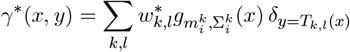

*and the entropic transport plan γ*^*ε*^ *converges to γ*^*^ *in the sense of the narrow convergence, i*.*e*., *for any bounded continuous functions ϕ* ∈ *C*_*b*_(ℝ^*n*^ × ℝ^*n*^),

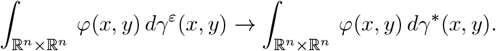

*Moreover*, 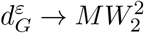 *as ε* → 0 *with the following estimate:*

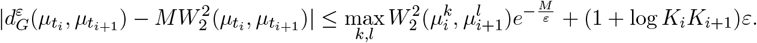

Theorem 3 enables us to go back and forth between the discrete and continuous optimal transports through the entropic transport plan (8). Thus, it provides a versatile trajectory inference framework (from continuous OT) with low computational complexity (from discrete OT) (Figs. S1–S2). Moreover, from the entropic transport plan (8), it is possible to make a map that transports the least cost from position *x* (entropic barycentric projection map) and the interpolated distribution between 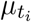 and 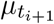 (entropic displacement interpolation). These mathematical results can be used to construct the functions of scEGOT, such as velocity fields of cell differentiation (cell velocity) and space-time continuous videos with gene expressions. Using cell velocity, it is also possible to infer the GRNs between adjacent time points and to reconstruct Waddington’s landscape in gene expression space (for more information on the theory of scEGOT’s functions, see *Methods* section).

### Biological application of scEGOT to human PGCLC induction system dataset

This section presents a biological application of scEGOT to time-series single-cell gene expression data and its validation. In particular, we consider the human primordial germ cell-like cell (PGCLC) induction system in vitro. In this in vitro culture system, previous studies have shown that genes such as *EOMES, GATA3, SOX17, TFAP2C* are critical genes for PGCLC differentiation [37, 38, 39, 40, 41]. However, the induction rate of PGCLC is only approximately 10 to 40 %, and the precise molecular mechanisms underlying the PGCLC differentiation and PGCLC/somatic cell segregation remain poorly understood. Using the functions of scEGOT, we characterize the progenitor cell population and elucidate the molecular mechanism of PGCLC/somatic cell segregation.

### Clustering

scEGOT allows for the manual division of clusters. From a biological perspective, we set the number of clusters *K*_*i*_ as 1, 2, 4, 5, and 5 for day 0, day 0.5, day 1, day 1.5, and day 2, respectively (Figs. 3A and S6A). It is also possible to determine the number of clusters using information criterion methods such as AIC or BIC. Each cluster is characterized by the high expression of key lineage-specific markers, enabling us to identify distinct lineages accurately. Specifically, we identified PGCLC (*NANOG*^+^, *SOX17* ^+^, *TFAP2C* ^+^, *PRDM1* ^+^), amnion-like cell (AMLC) (*TFAP2A*^+^, *TFAP2C* ^+^, *GATA3* ^+^), endoderm-like cell (EDLC) (*GATA6* ^+^, *SOX17* ^+^, *FOXA2* ^+^), and extra-embryonic mesenchyme-like cell (EXMCLC) (*HAND1* ^+^, *FOXF1* ^+^) (Fig. 3B and Figs. S6B–C). Identifying five clusters on days 1.5 and 2 reflects the differentiation states of the ExMCLC, which are divided into two clusters due to differences in their maturation stages. These cluster annotations are consistent with previous studies [42, 43, 44].

**Figure 3:**
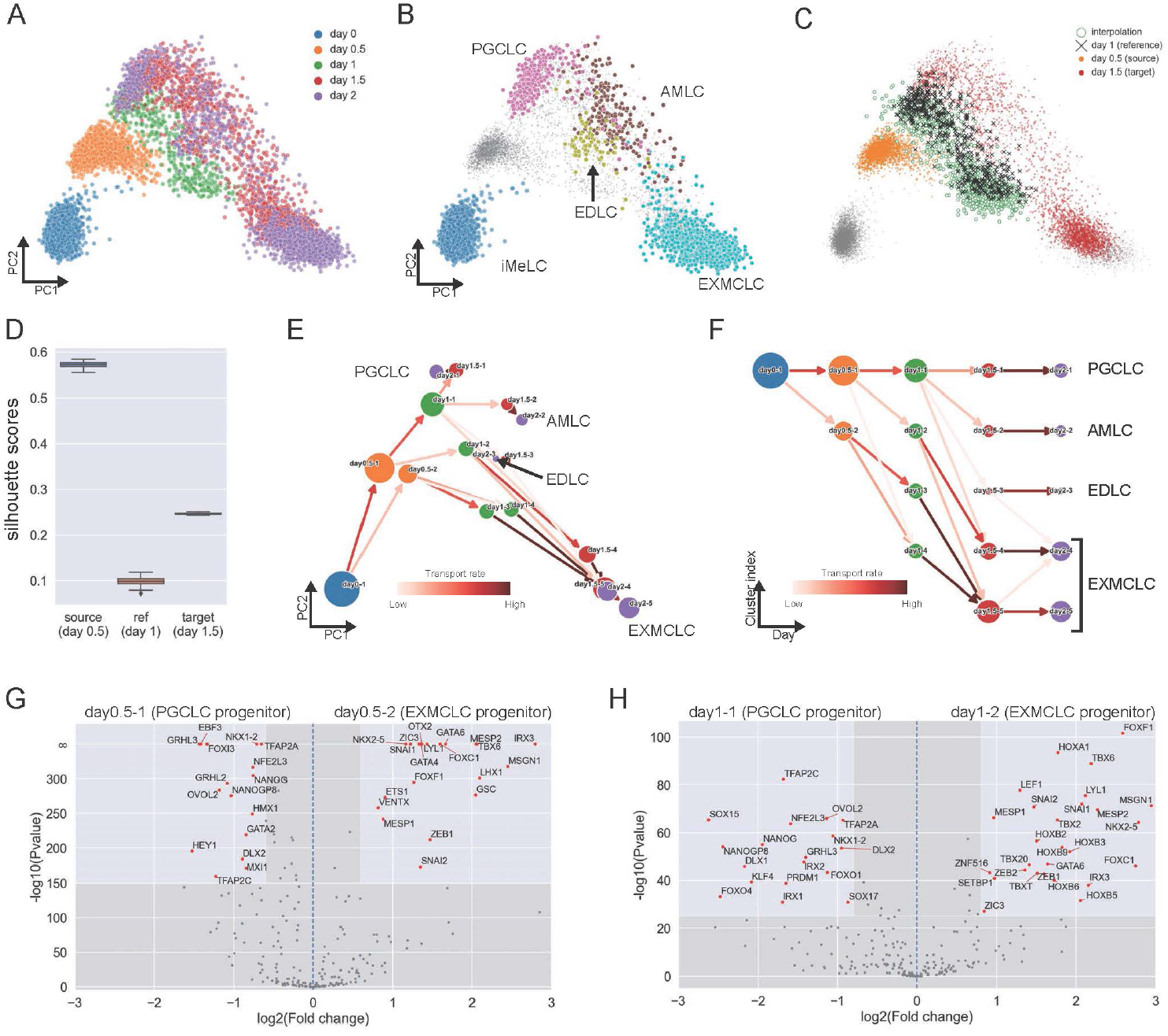
Application of scEGOT to the human PGCLC induction system dataset and identification of differentiation pathways. (A), (B) PCA plots of the PGCLC induction dataset. The cells are colored according to (A) experimental day and (B) cell type. The gray points in (B) are cells at the middle stages (days 0.5–1.5). (C) Verification of scEGOT interpolation. Comparison between the reference distribution (day 1) and scEGOT interpolation distribution generated by datasets at days 0.5 and 1.5 on PCA coordinates. (D) Box plot of the silhouette scores over 100 trials of the scEGOT interpolation versus the source (day 0.5)/reference (day 1)/target (day 1.5). (E), (F) Cell state graphs on PCA coordinates and by a hierarchical layout. The colors of the edges denote the transport rates 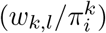 (G), (H) Volcano plots for day 0.5 (day0.5-1 and day0.5-2) and day 1 (day1-1 and day1-2) clusters. The horizontal and vertical lines show the log2 fold change, where the fold change indicates the rate of variation of gene expression and the negative common logarithm of *p*-values calculated from the independent samples t-test, respectively. The annotated points show cluster-specific genes with (G) | log_2_(Fold change)| *>* 0.6 and − log_10_(Pvalue) *>* 150 and (H) | log_2_(Fold change)| *>* 0.8 and − log_10_(Pvalue) *>* 25.

### Verification of scEGOT interpolation

We verify the accuracy of the entropic displacement interpolation of scEGOT. We set the data at day 1 as the reference distribution and generated the interpolation distribution using the data at day 0.5 (two clusters) and 1.5 (five clusters). The analysis shows that the interpolation distribution (1,000 cells) properly reproduces the reference distribution (Fig. 3C). This was quantitatively verified using the silhouette score (Fig. 3D), showing that the interpolation by scEGOT is well overlapped with the reference data rather than the source and target data. Here, the silhouette score indicates that the clusters are separated (overlapped, resp.) if the value is close to one (zero, resp.). We also performed the same verification for all other days and calculated the silhouette score (Figs. S3 and S4).

### Trajectory inference with cell state graph

In this section, we analyze the trajectory of cell differentiation in the human PGCLC induction system using the cell state graph (Fig. 3E) generated by scEGOT and gain molecular insights into the mechanism that segregates PGCLC from somatic lineages. We also applied a standard trajectory inference method, PAGA [13], to the same dataset (Fig. S5). The PAGA graph represents cell clusters and their similarities using nodes and edges. However, in datasets with rapidly changing cell states, such as transitions from iMeLC to day 1, the PAGA graph edges, determined by distances between clusters, may fail to capture developmental connectivity accurately. In contrast, the cell state graph generated by scEGOT successfully captures cell differentiation developments across substantial shifts in cell states. It enables us to identify key transition points and understand the temporal sequence of differentiation events. This feature makes the cell state graph more informative in terms of capturing the dynamic process of differentiation, providing clearer insights into how cell states evolve over time.

The cell state graph shows four primary differentiation pathways leading to PGCLC, AMLC,

EDLC, and EXMCLC as follows:

PGCLC: day0-1 → day0.5-1 → day1-1 → day1.5-1 → day2-1;

AMLC: day0-1 → day0.5-1 → day1-1 → day1.5-2 → day2-2;

EDLC: day0-1 → day0.5-1 → day1-1 → day1.5-3 → day2-3;

EXMCLC: day0-1 → day0.5-2 → day1-3 → day1.5-5 → day2-5.

The top path in the cell state graph (Fig. 3F) represents the PGCLC differentiation pathway. This path features key PGCLC markers such as *NANOG, SOX17, TFAP2C* and *PRDM1* [37, 38, 40, 41] (Fig. S6C). The expression levels of these markers increase from day 0.5 to day 1.5. It is of note that major EXMCLC progenitors are segregated as early as day 0.5 (see *Space-time continuous gene expression analysis* section for further analysis). Additionally, EXMCLC can be generated through alternative pathways, most typically, day0-1 → day0.5-1 → day1-2 → day1.5-4 → day2-4, suggesting that day0.5-1 cells retain a competence to differentiate into EXMCLC. Indeed, while there are pathways from the PGCLC progenitors until day 1 to the somatic cell populations, there is no pathway in the opposite direction. This result is reminiscent of Weismann’s barrier [45].

To gain a deeper understanding of PGCLC/EXMCLC segregation, we examine the difference in the gene expression value between the clusters. The volcano plots (Figs. 3G–H) show the comparison between two branched clusters at day 0.5 (day0.5-1 and day0.5-2) and day 1 (day1-1 and day1-2) of the PGCLC and EXMCLC lineages. Notably, we found that *TFAP2A* is expressed in the PGCLC progenitor cell population (day0.5-1). Previous studies have shown that this gene plays an important role in PGCLC differentiation [46, 42, 47], suggesting that the cell state graph generated from scEGOT has the ability to capture the progenitor cell population of PGCLC differentiation.

We also discovered that *NKX1-2* is highly expressed in the PGCLC progenitor compared to the somatic cells. This gene is known to be expressed during mesoderm development [48]. However, its role in PGCLC differentiation has been unknown. This finding will be further investigated in later sections using other functions of scEGOT.

Conversely, genes that are highly expressed in the EXMCLC pathway may also play a critical role in PGCLC/EXMCLC segregation. The early mesoderm markers (*MESP1, MESP2*) and the later mesoderm markers (*GATA6, FOXF1*) are significantly upregulated on day0.5-2. This shows that these somatic genes might act as repressors of PGCLC during the segregation.

Furthermore, when comparing the clusters day1-1 and day1-2, the former, which is on the way to the PGCLC pathway, shows enrichment of key PGCLC specification genes, namely *NANOG, TFAP2C, SOX17, KLF4, SOX15* and *PRDM1* (Fig. 3H). This indicates that the PGCLC path is more specified on day 1.

In mouse PGC, genes such as *stella* and *fragilis* are actively expressed, while HOX genes such as *Hoxa1, Hoxb1* are repressed [49, 50, 51, 52]. On the other hand, we did not observe any HOX gene expression in human PGCLC. Interestingly, in our induction system, *HOXA1, HOXB2, HOXB3, HOXB5* and *HOXB6* are highly expressed in the EXMCLC lineage.

From the above observation, the cell state graph captures the trajectories of human PGCLC induction well and can potentially find key genes by analyzing differentially expressed genes among the clusters. In particular, the cell state graph succeeded in tracing PGCLC/somatic cell segregation and identified key genes that may enhance or suppress PGCLC differentiation, such as *NKX1-2, MESP1, MESP2*, and *GATA6*.

### Velocity analysis

We investigated the dynamics of the developmental process in human PGCLC induction using the cell velocity generated by scEGOT and compared our method with the RNA velocity [54, 55]. RNA velocity is a standard method to estimate the velocity field in the gene expression space from scRNA-seq data, and it is also widely used to infer cell trajectories [56].

The comparison of the normalized speeds between cell velocity and RNA velocity showed that, despite some differences in the iMeLC and early mesoderm populations, the overall behavior of cell and RNA velocities is consistent between the two methods (Fig. S7). This consistency suggests that both velocity fields accurately reflect the dynamics of lineage determination, aligning with biological observations (Figs. 4A–B).

**Figure 4:**
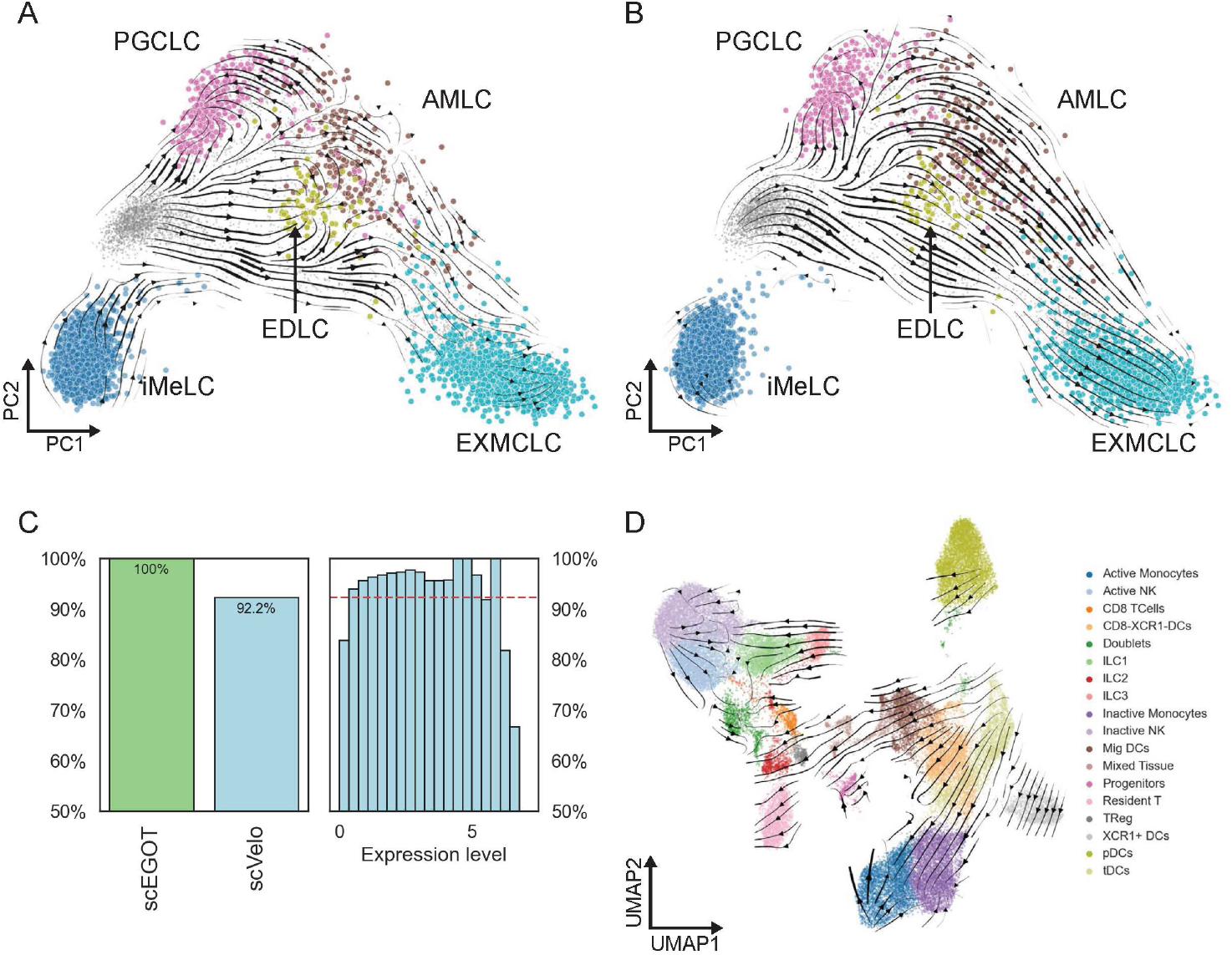
Comparison of velocities between scEGOT (cell velocity) and scVelo (RNA velocity). (A) Stream plot of cell velocity generated by scEGOT. (B) Stream plot of RNA velocity generated by scVelo (stochastic mode). (C) Left: Percentage of genes for which velocities can be computed by scEGOT and scVelo. Right: Histogram of the coverage of scVelo for mean expression levels. The red dashed line is the total coverage of scVelo (92.3%). (D) Cell velocity for scATAC-seq data of mouse innate immune cells at three-time points (days 0, 1, and 28) [53].

Furthermore, scEGOT can provide the velocities for all the genes, whereas the RNA velocity cannot calculate the velocities for genes without a sufficient amount of detection of unspliced RNA [54, 55] (Fig. 4C). We also emphasize that since the cell velocity is a data-driven method, it can be applied not only to scRNA-seq data but also to any other time-series single-cell data, such as scATAC-seq data. We apply it to time-series scATAC-seq data for innate immune cells from mouse-draining lymph nodes (Fig. 4D). The flow of NK cells and monocytes from inactive to active states can be observed. Overall, the cell velocity allows us to perform a comprehensive velocity analysis.

### Space-time continuous gene expression analysis

To study the cell differentiation dynamics, such as the bifurcation time of PGCLC/somatic cell segregation, we constructed the time interpolations of cell populations and the time-continuous gene expression dynamics (animation) using the entropic displacement interpolation (Fig. 5 and Movies S1–S4). The result clearly shows the temporal evolution of cell differentiation and gene expression patterns for the marker genes of PGCLC. In particular, as early as 0.25 and more clearly at 0.75 days, the clusters *NKX1-2* ^+^ and *NKX1-2* ^−^ are separated, and the former cluster exactly moves to the PGCLC population (*TFAP2C* ^+^, *PRDM1* ^+^) (Movies S1, S3).

**Figure 5:**
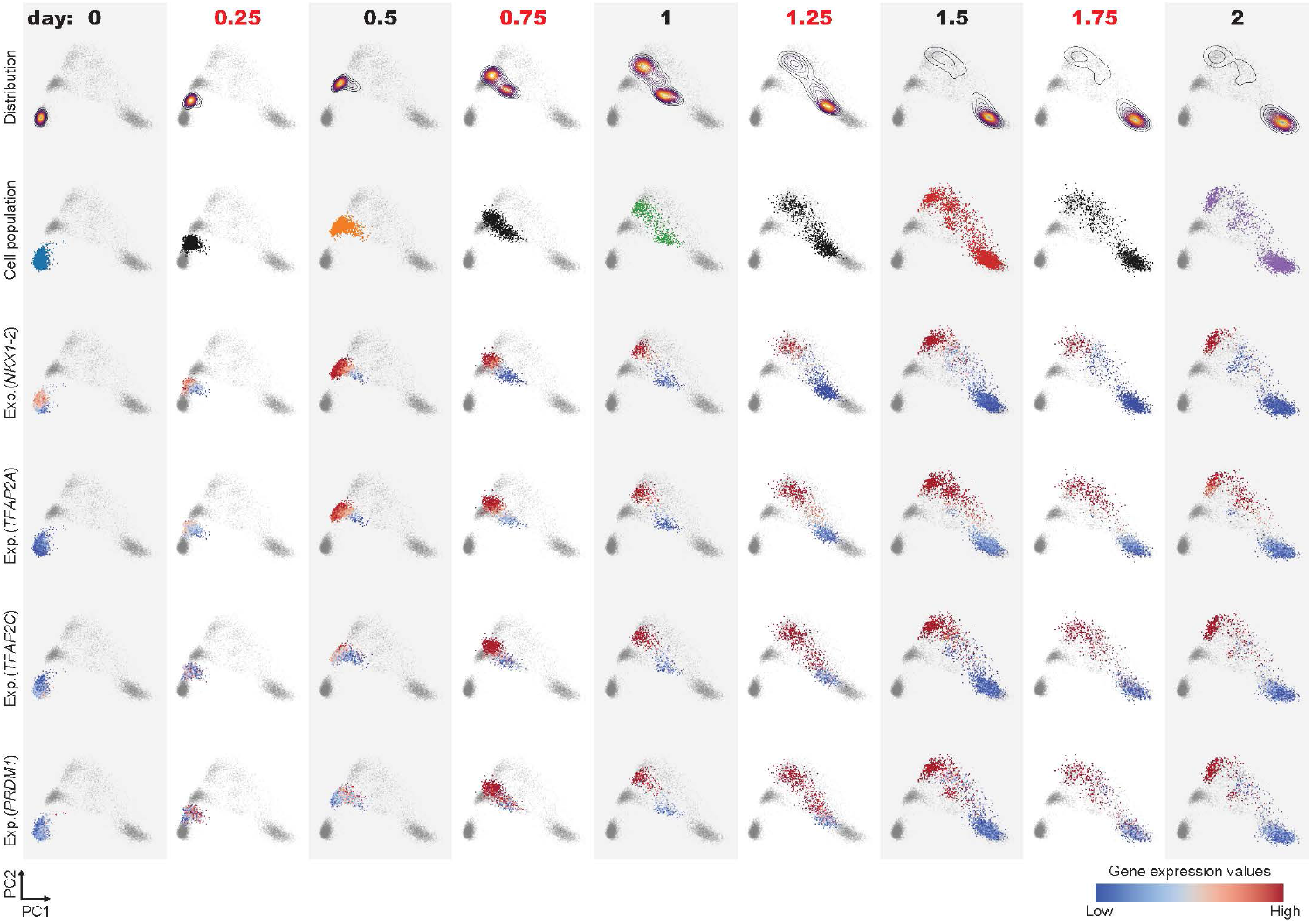
Input scRNA-seq data (gray columns) and interpolated data (white columns) on the top two principal components. The first row shows the contour plots of the Gaussian mixture distributions. The second row denotes the cell populations of the real scRNA-seq data (gray columns) and those generated by the interpolated Gaussian mixture distributions (white columns). The third to sixth rows show the gene expression values of *NKX1-2, TFAP2A, TFAP2C*, and *PRDM1* in the cell populations.

On the other hand, although the cluster *TFAP2A*^+^, which has been reported as the progenitor cell population of the PGCLC in previous studies [46, 42, 47], shows a similar tendency to the cluster *NKX1-2* ^+^ until day 1, *TFAP2A* is also highly expressed in the AMLC population after this day, implying that *TFAP2A* alone cannot identify the PGCLC progenitor cell population. This analysis suggests that *NKX1-2* is one of the earliest marker genes of the PGCLC progenitor cell population, and its expression guides the cell population to the PGCLC pathway.

It also shows that the bifurcation of PGCLC/somatic cell segregation occurs much earlier, as early as 0.25 days.

### GRN analysis

We inferred the GRNs for PGCLC induction. We computed the GRN matrix using the cells in the clusters on the PGCLC pathway, which is determined by the cell state graph analysis. We then extracted a subnetwork of the GRN specified by the key genes of the human PGCLC specification *PRDM1, SOX17, TFAP2C* and two candidates *TFAP2A* and *NKX1-2* of PGCLC progenitor marker genes (Fig. 6A).

**Figure 6:**
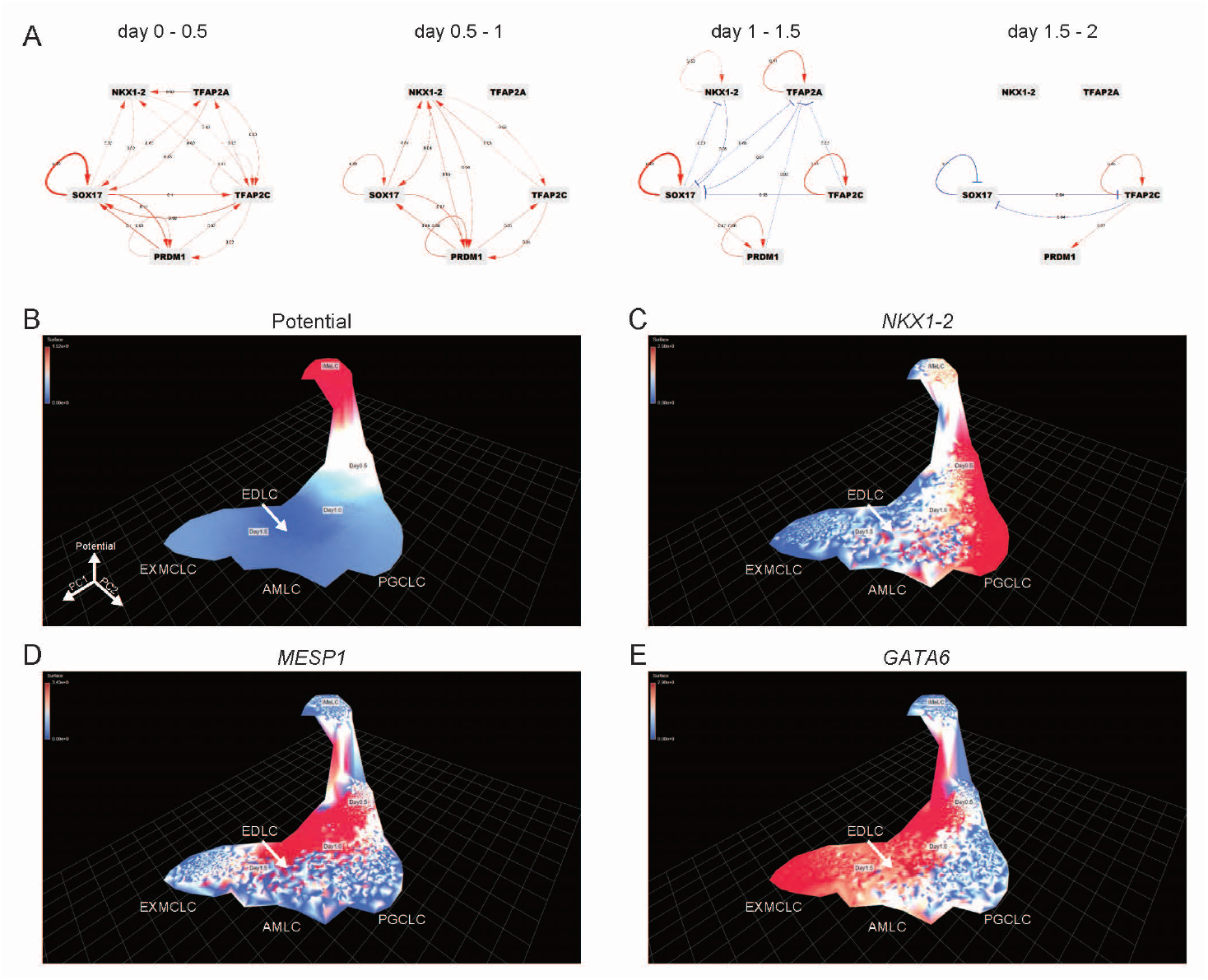
Reconstruction of Waddington’s landscape and inferring the GRNs for the human PGCLC induction system. (A) Gene regulatory networks of the human PGCLC induction system generated from scRNA-seq data at days 0–0.5, days 0.5–1, days 1–1.5, and days 1.5–2. (B)–(E) Reconstruction of Waddington’s landscape of human PGCLC induction data. The *x*-, *y*-, and *z*-axes denote the PC1, PC2 coordinates, and the Waddington potential, respectively. The visualization was prepared using CellMapViewer: https://github.com/yusuke-imoto-lab/CellMapViewer. The colors indicate (B) the magnitude of the potential, (C) *NKX1-2*, (D) *MESP1*, and (E) *GATA6* expression values.

On days 0–0.5, we found that both *TFAP2A* and *NKX1-2* activate *SOX17* which is essential for PGCLC differentiation [37, 38]. However, on days 0.5–1, *NKX1-2* continues to activate the other PGCLC regulators, whereas *TFAP2A* does not interact with them. This result suggests that *NKX1-2* may play a more essential role than *TFAP2A* during human PGCLC differentiation.

### Reconstruction of Waddington’s landscape

Finally, we reconstructed Waddington’s landscape to validate the differentiation ability of cells in the PGCLC induction system by scEGOT. Figs. 6B–E show the reconstructed Waddington’s landscape together with the expression values of the genes *NKX1-2, MESP1*, and *GATA6*.

The landscape (Fig. 6B) shows that the iMeLCs, which are characterized by high levels of pluripotent genes (*SOX2, NANOG, POU5F1*) and relatively low levels of early mesoderm genes (*TBXT, MIXL1, EOMES*) (Fig. S6B), are located at the top of the potential. This suggests that these cells have high plasticity, which is consistent with biological knowledge.

In addition, viewing the gene expression values on the landscape allows us to understand the contribution of the genes to cell differentiation. For instance, it can be seen that *NKX1-2* contributes to the early stage of PGCLC (Fig. 6C). In contrast, *MESP1* and *GATA6* contribute to the early stage of EXMCLC and EDLC differentiation, respectively (Figs. 6D–E). Importantly, we emphasize that their expression levels are complementary at days 0.5-1, suggesting that these genes may play the role of landscape pegs, forming a barrier between the PGCLC and the somatic cell pathways.

## Discussion and conclusions

We have formulated an entropic Gaussian mixture optimal transport (EGOT) and proved the mathematical properties of EGOT. Based on EGOT, we have also proposed a novel single-cell trajectory inference framework (scEGOT) that infers cell differentiation pathways and gene expression dynamics from time-series scRNA-seq data. Furthermore, we applied scEGOT to the human PGCLC induction system and validated it biologically.

scEGOT provided us with the following outputs in biological applications. The cell state graph, which represents the cell clusters and their transport rates as a graph model, showed the trajectories between the cell populations. The cell velocity generated by the entropic barycentric projection map has expressed the cell differentiation flow, which describes more detailed and consistent structures in the case of human PGCLC than the conventional method. The entropic displacement interpolation, which can generate a pseudo-cell population at any intermediate time, has simulated the temporal evolution of gene expression dynamics in single-cell resolution and has certainly identified the bifurcation time. The GRN analysis has contributed to the discovery of candidates of upstream genes that induce a cell differentiation system. The reconstruction of Waddington’s landscape, constructed based on the potential of the gradient flow associated with the cell velocity, has summarized the pathway of cell differentiation and represented the cell potency of each cell type. We note that all the outputs represented by the PCA coordinates can be replaced by those with other coordinates generated by other dimensionality reduction methods, such as uniform manifold approximation and projection (UMAP) [57], which provides biological interpretations from different aspects (Fig. S8).

Through the abovementioned scEGOT analysis, we have identified genes such as *NKX1-2, MESP1*, and *GATA6* as potential regulators for human PGCLC differentiation or its divergence towards somatic lineages. The biological functions of these genes in human PGCLC are still unknown and are expected to be revealed through biological experiments.

Looking back on this paper, we have revisited an epigenetic world that was once envisioned by Waddington. He viewed the process of cell development as a canal and stated that a cell within a region denoting a particular state would be carried through the canals to a single specified cell, which is the corresponding steady state. He modeled such a time transition as a phase-space diagram of development (Fig. 3 in [1]), which is a similar concept to the cell state graph. He then modeled the epigenetic landscape, which is a well-known potential model, by simply describing the phase-space diagram in a three-dimensional picture (Fig. 4 in [1]). He further explained that the chemical states of the genes and their regulatory system underlie the landscape model (Fig. 5 in [1]).

Overall, scEGOT has realized the images envisioned by Waddington through a comprehensive mathematical framework called EGOT and has succeeded in capturing complex cell differentiation systems and their dynamics by analyzing the cell state graph, cell velocity, interpolation analysis, gene regulatory network, and reconstruction of the epigenetic landscape.

We believe that scEGOT is useful for a broader range of biological data because it is a data-driven method. In other words, all scEGOT settings are never limited to scRNA-seq data. For example, scATAC-seq data containing open chromatin information could be applied in scEGOT to provide a deeper explanation of cell differentiation and lineage fate determination at epigenome resolution (Fig. 4D). Indeed, the epigenetic regulations during cell differentiation, such as the regulations of cis-regulatory elements and transposon elements, are fundamental questions, and our approach will be instructive in addressing such questions.

## Methods

In what follows, we describe the theory of the functions of scEGOT.

### Cell state graph by EGOT

In the case of point clouds obtained from the time-series scRNA-seq data, each Gaussian distribution and its weight can be regarded as a certain cell type and its existence probability, respectively. Then, the solution of EGOT, which represents how much weight is transported between the Gaussian distributions at adjacent time points, can be interpreted as the degree of relationship between these cell types.

Based on this interpretation, we define a cell state graph of cell differentiation as follows. For each adjacent times *t*_*i*_, *t*_*i*+1_, a complete weighted bipartite graph (*V*_*i*_, *W*_*i*_) in ℝ^*n*^ is constructed as

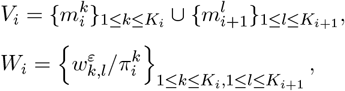

where the set *V*_*i*_ of nodes consists of the mean vectors at times *t*_*i*_ and *t*_*i*+1_ representing the locations of the cell types, and the set *W*_*i*_ of the weighted edges are given by the normalized solution of the EGOT (called transport rate) corresponding to the degree of relationship of the cell types. Then, the cell state graph (*V, W*) is constructed by combining (*V*_*i*_, *W*_*i*_) for all the time stages. From this cell state graph, we can study the state transitions of the cell population in the temporal evolution.

### Entropic barycentric projection map and cell velocity

The entropic transport plan *γ*^*ε*^(*x, y*) can be regarded as the weights to be transported from the point *x* to *y*. By averaging over *y*, we can define an entropic barycentric projection map of the EGOT, which transports the least cost from position *x*, as follows:

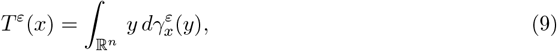

where 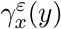 is the disintegration of the entropic transport plan *γ*^*ε*^ with respect to the first marginal 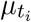, i.e., 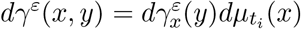 Then, by (8), we can compute the entropic barycentric projection map *T*^*ε*^ explicitly.

#### Proposition 4.

*The entropic barycentric projection map with respect to γ*^*ε*^ *is expressed as*

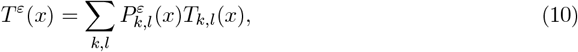

*where* 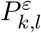 *denotes*

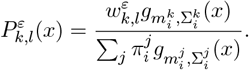

From the definition, the entropic barycentric projection map *T*^*ε*^ represents where a cell 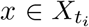 at time *t*_*i*_ moves at time *t*_*i*+1_ in the gene expression space. Accordingly, we define the rate of change of the gene expression for a cell *x* from the time *t*_*i*_ to *t*_*i*+1_, called the cell velocity, as

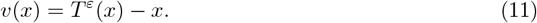

The cell velocity *v*(*x*) expresses which direction and how much the gene expression of the cell *x* changes. A high speed |*v*| implies a significant change of the cell in cell differentiation, whereas a low speed |*v*| indicates that the gene expression hardly changes and is close to the steady state.

### Entropic displacement interpolation and gene expression animation

One of the great advantages of the optimal transport theory is that we can obtain the displacement interpolations between the probability distributions, allowing us to study the dynamics of cell differentiation. We define the following entropic displacement interpolation between 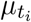 and 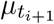

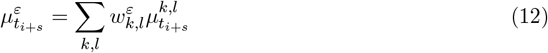

at *t*_*i*+*s*_ = (1 − *s*)*t*_*i*_ + *st*_*i*+1_, *s* ∈ [0, 1]. Here 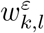 is the solution (3), and 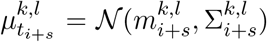 with 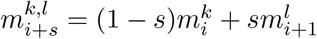 and

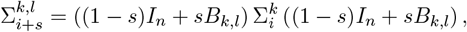

where *I*_*n*_ is the *n* × *n* identity matrix and

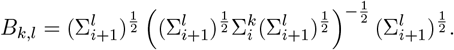

#### Theorem 5.

*The probability distribution*

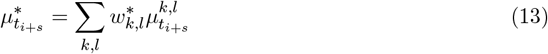

*is the geodesic with the maximum entropy between* 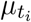 *and* 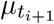 *in the space G*_2*n*_(∞) *equipped with the distance MW*_2_, *where w*^*^ *is the solution* (6). *Moreover, for any s* ∈ [0, 1], *the entropic displacement interpolation* 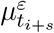 *converges to* 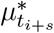 *narrowly*.

From Theorem 5, we see that the entropic displacement interpolation (12) is an approximation of the geodesic (13) in the space *G*_2*n*_(∞) equipped with the distance *MW*_2_. Through this entropic displacement interpolation (12), we can generate an interpolation distribution between 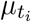 And 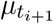, which means that the interpolation of the adjacent scRNA-seq data 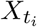 and 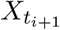 can be generated. Therefore, by successively constructing the interpolations for all the time stages, we can create an animation of the gene expressions to track the time-varying dynamics of cell differentiation.

### Estimation of GRNs

Once the cell velocity (11) representing the changes in the gene expressions is obtained, it can be used to estimate the gene regulatory network (GRN) that drives the cell differentiation dynamics. Let 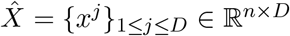 be a cell population consisting of *D* cells for which the GRN is to be obtained. We assume that the gene expressions in the cell population 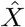 are driven by linear dynamics

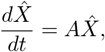

where *A* ∈ ℝ^*n*×*n*^ is the matrix characterizing the GRN and represents the effect of the expression level of each gene on the expression dynamics of the other genes. By the cell velocity (11), we may assume that 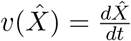. Thus, we obtain

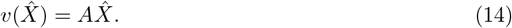

Based on (14), the GRN matrix *A*^*^ is estimated by solving the following regression problem:

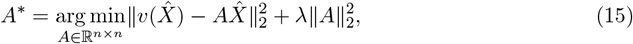

where *λ >* 0 is a regularization parameter. In terms of each cell *x*^*j*^, this can be expressed as

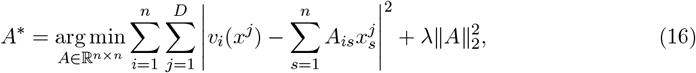

where *v*_*i*_(*x*^*j*^) is the *i*-th component of *v*(*x*^*j*^). By solving the problem (15) or (16) at each adjacent time point, we can estimate the GRNs driving cell differentiation.

We compared our approach to other methods for GRN inference using the BEELINE bench-mark framework and its test data [58]. SCODE [59] is an algorithm similar to our method included in BEELINE. While both scEGOT and SCODE are designed to infer GRNs from scRNA-seq data, they differ in their underlying methodologies. SCODE utilizes ordinary differential equations (ODEs) to model gene expression dynamics and focuses on reconstructing the regulatory networks using linear ODEs and linear regression. SCODE applies this approach to pseudotime data, treating it as a time-like variable to infer GRNs from scRNA-seq data. In contrast, scEGOT can construct GRNs that capture dynamic regulatory interactions over time by using cell velocity. This enables scEGOT to directly incorporate the continuous temporal dynamics of cell differentiation, providing a more precise representation of gene regulation over time.

When applying scEGOT to the BEELINE benchmark test data, we first clustered the data based on pseudotime and then treated these clusters as virtual time-series scRNA-seq data. This approach allowed us to apply scEGOT’s GRN inference algorithm to single-time scRNA-seq data. We found comparable performance in terms of GRN inference (Fig. S9). We also note that the strength of scEGOT’s GRN inference lies in its ability to construct time-dependent (dynamic) GRNs using the cell state graph, which goes beyond static GRN inference typically used for single time point data. This dynamic approach provides a more comprehensive view of the regulatory interactions throughout the differentiation process, allowing scEGOT to offer unique insights into gene regulation over time.

### Construction of Waddington’s landscape

From the cell velocity (11), we can construct Waddington’s landscape in the gene expression space. The Helmholtz–Hodge decomposition implies that the cell velocity can be written as

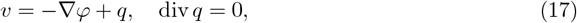

where *ϕ* is the gradient potential and *q* is the divergence-free part. Since Waddington’s landscape is a gradient system, the potential *ϕ* can be regarded as the realization of Waddington’s landscape in the gene expression space. To construct *ϕ*, we take the divergence operator in the equation (17) and obtain the following differential equation

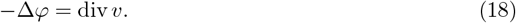

The partial differential equation (18) is not uniquely solvable in general since the boundary conditions are undefined. In the following, instead of solving (18) directly, we consider it as an equation on the *k*-nearest neighbor graph of cells and look for its least-squares and least-norm solution. In other words, we consider the potential *ϕ* as the solution to the following minimization problem:

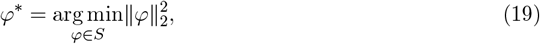

where *S* is the set of solutions to the least-squares minimization problem

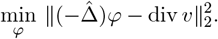

Here, 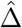 denotes the graph Laplacian on the *k*-nearest neighbor graph of the cells. The solution of the minimization problem (19) can be written as

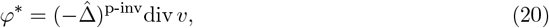

where 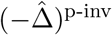 denotes the Moore–Penrose pseudoinverse of (− 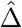). Note that the divergence of the cell velocity *v* on the right-hand side in (20) can also be computed concretely, as the cell velocity (11) is obtained explicitly. Thus, the following theorem holds.

#### Theorem 6.

*The potential ϕ*^*^ *is represented by*

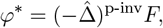

*With*

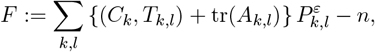

*where* (·, ·) *denotes the standard Euclidean inner product of vectors and C*_*k*_ *denotes*

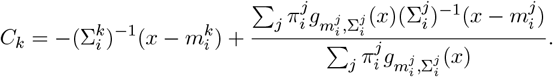

Through this procedure, we can construct the gradient potential (20) of the cells as in Waddington’s landscape.

We compared the Waddington potential reconstructed by scEGOT with the potential inferred by CytoTrace2 [60], which represents the potency score. In CytoTrace2, the potency score ranges from 0 to 1, but the values appear to be reversed in the PGCLC dataset. For example, iMeLC has a potency score closer to 0, while other cell types show scores closer to 1. This is opposite to the expected biological knowledge, where iMeLC should have a higher score. However, when we consider (1-the potency score), we found that it closely matches the Waddington potential from scEGOT. This suggests that both methods capture a similar Waddington landscape (Fig. S10).

### Culture of human iPSC

We are utilizing the 585B1 BTAG hiPSC (46, XY) male cell line, which is maintained in StemFit AK02 medium (Ajinomoto, Tokyo, Japan) on iMatrix-511 (Nippi, Tokyo, Japan) coated cell culture plates. For cell passage, cells are dissociated using 0.5x TrypLE Select, a mixture of TrypLE Select (Gibco, 12563-011) and PBS(-) (Phosphate Buffered Saline, Nacalai Tesque, 11480-35) containing 0.5 mM EDTA (Nacalai Tesque, 06894-14), at 37^°^C for 10 minutes. The cells are resuspended in AK02 medium supplemented with 10 M Y27632 (Tocris, 1254) and replaced with fresh medium without Y27632 after 24 hours. The medium is changed every other day.

### hPGCLC induction

The methods for hPGCLC induction were carried out as previously described (Sasaki et al., 2015). hiPSC were induced to iMeLC in fibronectin-coated (Millipore, FC010) 12-well plates using induction medium [GMEM (Gibco, 11710-035) containing 15% KnockOut Serum Replacement (KSR) (Gibco, 10828-028), 1% MEM Non-Essential Amino Acids Solution (NEAA) (Gibco, 11140-050), 1% penicillin-streptomycin, 2 mM L-glutamine (Gibco, 25030-081), 2 mM sodium pyruvate (Gibco, 11360-070), and 0.1 mM 2-mercaptoethanol], supplemented with 3 M CHIR99021 (Tocris, 4423), 50 ng/ml activin A (PeproTech, AF-120-14), and 10 M Y27632. After 42-48 hours, iMeLC were dissociated into single cells using 0.5x TrypLE Select for 10 minutes at 37^°^C and transferred to a low-cell-binding V-bottom 96-well plate (Greiner, 651970) containing hPGCLC induction medium [GMEM with 15% KSR, 1% NEAA, 1% penicillin-streptomycin, 2 mM L-glutamine, 2 mM sodium pyruvate, and 0.1 mM 2-mercaptoethanol], supplemented with 200 ng/ml BMP4 (R&D Systems, 314-BP), 100 ng/ml SCF (R&D Systems, 255-SC), 10 ng/ml LIF (Merck Millipore, LIF1010), 50 ng/ml EGF (PeproTech, AF-100-15), and 10 M Y27632. The hPGCLC induction medium was not changed until analysis.

### 10X experiment and dataset

The samples for scRNA-seq analysis were collected at iMeLC for 46 hours. At PGCLC aggregate days 0.5, 1, 1.5, and 2, iMeLCs were dissociated as previously described. PGCLC aggregates were washed with PBS(-), then dissociated using 0.25% TrypsinEDTA for 1015 minutes at 37^°^C, followed by gentle pipetting 10 times. The trypsin was neutralized with 10% FBS in DMEM, and the cells were washed once with 0.1% BSA. ScRNA-seq sample preparation and library construction for 10X data were performed using the 10X Genomics Chromium Controller (10X Genomics) and Chromium Single Cell 3’ Reagent Kits v3.1, following the manufacturer’s instructions. The dataset for scRNA-seq consists of five-time points (*I* = 5); we refer to iMeLC as day 0, whereas 3D-cultured aggregates are denoted as day 0.5, day 1, day 1.5, and day 2, respectively (Fig. 3A).

### Preprocessing and parameter settings

As a preprocessing step for scEGOT, we applied a noise reduction method RECODE [61] and selected the top 2,000 genes with the highest normalized variances (variances divided by means) of non-mitochondrial genes, followed by log scaling. Log scaling is more important than other preprocessing steps for transforming the distributions of cell populations to Gaussian distributions, as mRNA expression levels are known to follow a log-normal distribution [62]. Thus, this transformation is crucial for approximating a multivariate Gaussian distribution suitable for scEGOT. We then performed PCA and applied scEGOT to the top 150 principal components (their cumulative contribution rate was 93.67%). It should be noted that increasing these values allows the data to be represented in more detail, but it also increases the computational cost. We set the regularization parameter of EGOT to *ε* = 0.01. We remark that the convergence result (5) indicates that the solution of EGOT remains largely unaffected even when the parameter *ε* is changed. In the *GRN analysis* section, we cut the small regulatory edges with a weight less than 0.02. The regularization parameter *λ >* 0 was automatically determined by cross-validation. In *Reconstruction of Waddington*’*s landscape* section, we constructed the *k*-nearest neighbor graph of cells (*k* = 15) on the top two principal components to compute the graph Laplacian.

## Declarations

### Ethics approval and consent to participate

The experiments involving hPGCLCs induced from hiPSCs were approved by the Institutional Review Board of Kyoto University and were also performed in accordance with the guidelines of the Ministry of Education, Culture, Sports, Science, and Technology (MEXT) of Japan.

### Consent for publication

It does not include individual data.

### Availability of data and materials

ScRNA-seq data of the human PGCLC induction system are available in the Gene Expression Omnibus (GEO) under identification number GSE241287. The Python software scEGOT is available through an open-source package at https://github.com/yachimura-lab/scEGOT.

## Authors’ contributions

T.Y., Y.I., S.T., M.S., and Y.H. designed the study; T.Y. and H.W. performed the research; T.Y. developed the new analytic tools; T.Y., Y.I., and M.Y contributed to the development of the new analytic tools; T.Y. and H.W. analyzed the data; H.W., Y.K., and Y.Y. collected and set-up the scRNA-seq data; and T.Y., H.W., Y.I., M.S., and Y.H. wrote the paper.

## Acknowledgements

This work was supported by the World Premier International Research Center Initiative (WPI).

## Supporting information for

In the following, proofs of propositions and theorems in this paper are shown. In addition, figures and animations that could not be included in the paper are also given.

### Proofs of propositions and theorems

#### Proof of Proposition 1

Since the discrete entropy *H* is a strongly concave function, the objective function of (2) is a strongly convex function. Therefore, the optimization problem (2) has a unique optimal solution. Also, we consider the following Lagrangian associated with the problem (2):

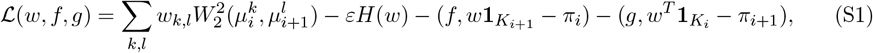

where 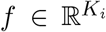 and 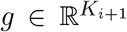 are the Lagrange multipliers and (·, ·) denotes the standard Euclidean inner product of vectors. The first-order condition of (S1) yields that

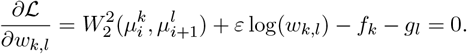

Thus the unique solution *w* = *w*^*ε*^ of (2) is given by

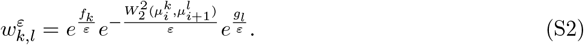

Putting 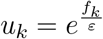 and 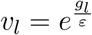 in (S2), then we obtain

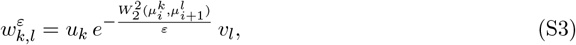

which is the desired conclusion.

#### Proof of Proposition 2

We consider the convergence of EGOT and the solution (S3) as *ε* → 0. By using an estimate similar to that for discrete entropic regularized optimal transports with Shannon entropy by Weed [1], we can show that the solution (S3) of EGOT exponentially converges to an optimal transport plan of 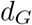 with the maximal entropy as *ε* → 0. That is, there exists *M >* 0 independent of *ε >* 0 such that

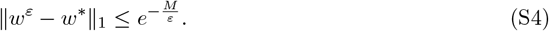

By using the estimate (S4), we have

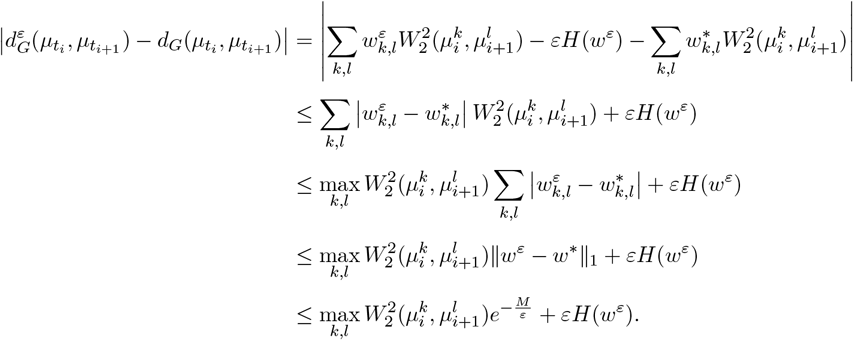

From Jensen’s inequality and the fact that the function − log *t* is convex, we obtain

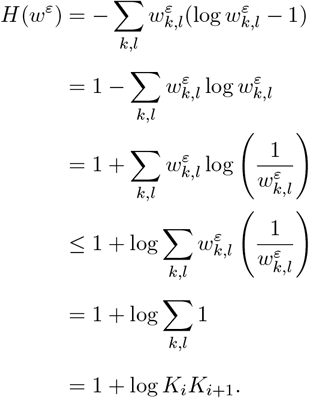

This leads to the following estimate

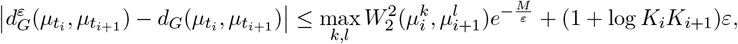

which is the desired conclusion.

#### Proof of Theorem 3

As shown in [2, Proposition 4, p.944], the discrete optimal transport problem

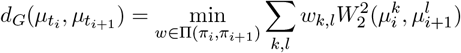

coincides with the continuous optimal transport problem (4). Therefore, by Proposition 2, we obtain the following estimate:

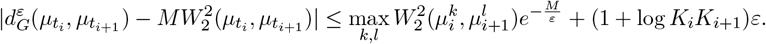

Moreover, for any bounded continuous functions *ϕ* ∈ *C*_*b*_(ℝ^*n*^ × ℝ^*n*^),

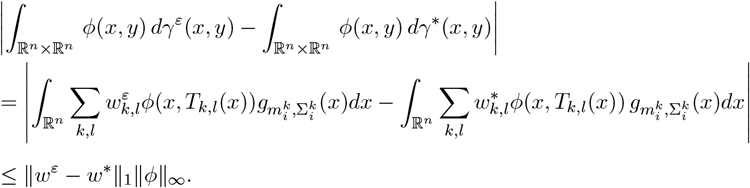

By using (S4), we obtain

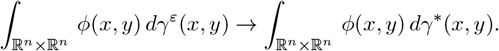

Thus, the entropic transport plan *γ*^*ε*^ converges to *γ*^*^ in the sense of the narrow convergence.

#### Proof of Proposition 4

Recall that the definition of the entropic barycentric projection map is

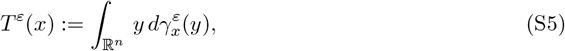

where 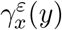 is the disintegration of the entropic transport plan *γ*^*ε*^ with respect to the first marginal 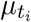, i.e.,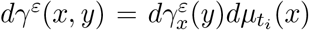. Then, by (8), the probability density of 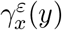 is given by

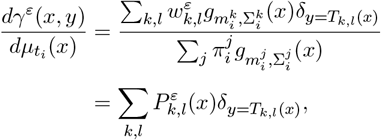

where 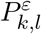 denotes

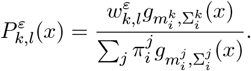

Thus, the entropic barycentric projection map (S5) can be computed as follows:

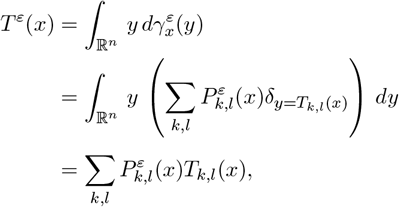

which is the desired conclusion.

#### Proof of Theorem 5

By the results of [2, Proposition 5 and Corollary 2, pp.946–948], the probability distribution

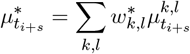

is the geodesic with the maximum entropy between 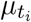 and 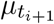 in the space *G*_2*n*_(∞) equipped with the distance *MW*_2_, where *w*^*^ is the solution of (6). Also, the proof that the entropic displacement interpolation 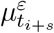 converges to 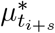 narrowly can be shown in the same way as Theorem 3. Indeed, for any bounded continuous functions *ϕ* ∈ *C*_*b*_(ℝ^*n*^),

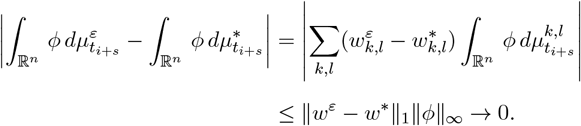

Here, we used (S4).

#### Proof of Theorem 6

##### Theorem 6

can be shown by direct calculation as follows:

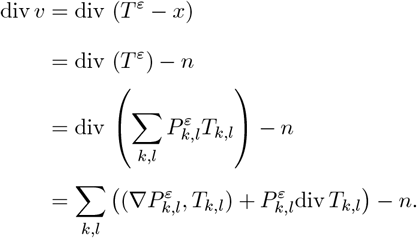

By the definition of 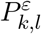, we have

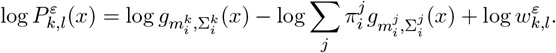

Then, we get

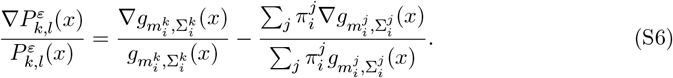

Also, in the same manner as above, by the definition of 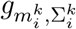 we have

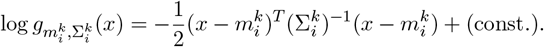

Then, we obtain

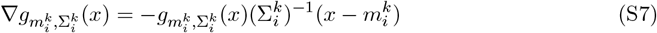

By substituting (S7) into (S6), we have

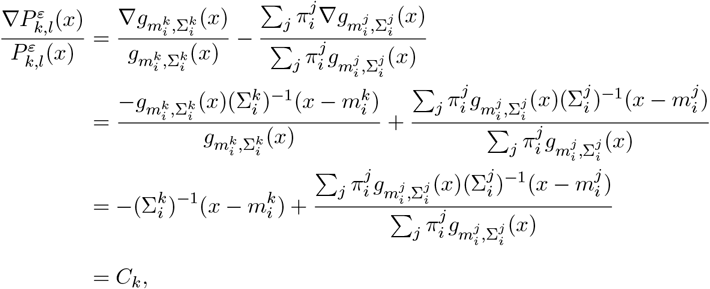

where we set *C*_*k*_ as the final expression. Therefore, we obtain

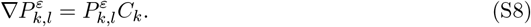

Also, let us recall that the optimal transport map between 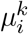 and 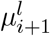 is given by

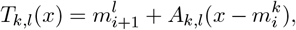

where *A*_*k,l*_ denotes

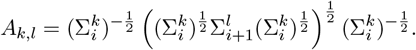

The direct computation implies that the following equality holds:

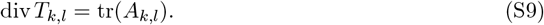

By (S8) and (S9), we obtain

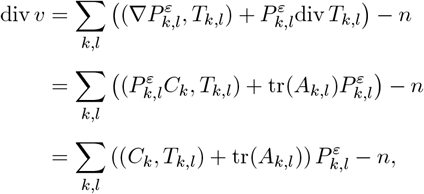

which is the desired conclusion.

## Figures

**Figure S1:**
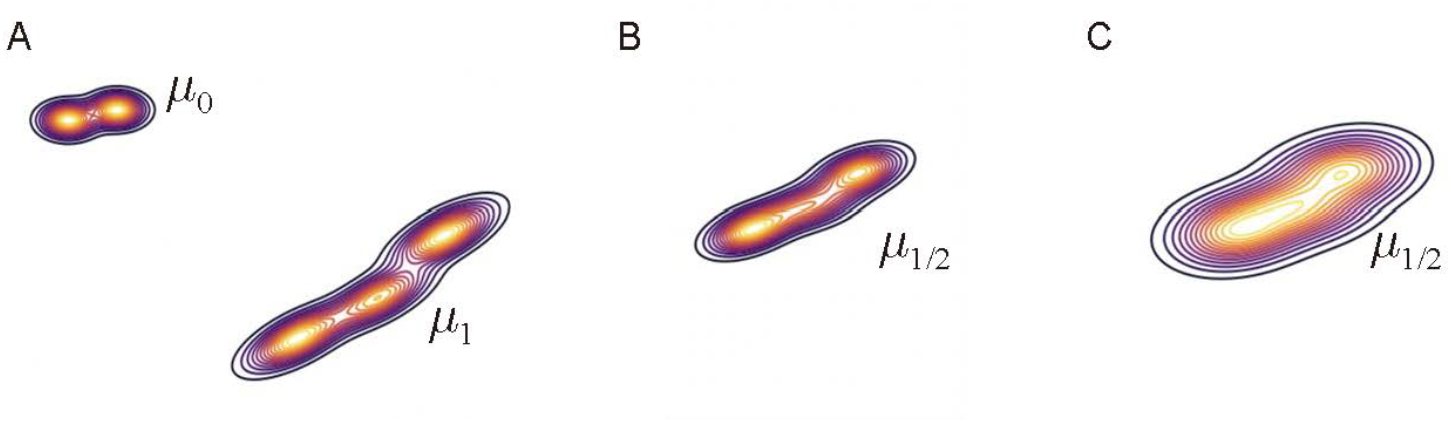
Comparison of computation time between interpolated distributions generated by EGOT and continuous OT. The experiments were performed on a MacBook Pro (14-inch, 2021) equipped with Apple M1 Max and 64GB RAM. (A) Source and target Gaussian mixture distributions *µ*_0_ and *µ*_1_ in 2D with two and three clusters, respectively. The space was divided into 40,000 segments. (B) Interpolation distribution *µ*_1*/*2_ generated by EGOT. The computation time was 41ms. (C) Interpolation distribution *µ*_1*/*2_ generated by using the POT library [3], specifically the ot.bregman.barycenter module. The computation time was 1h 44min 49s.

**Figure S2:**
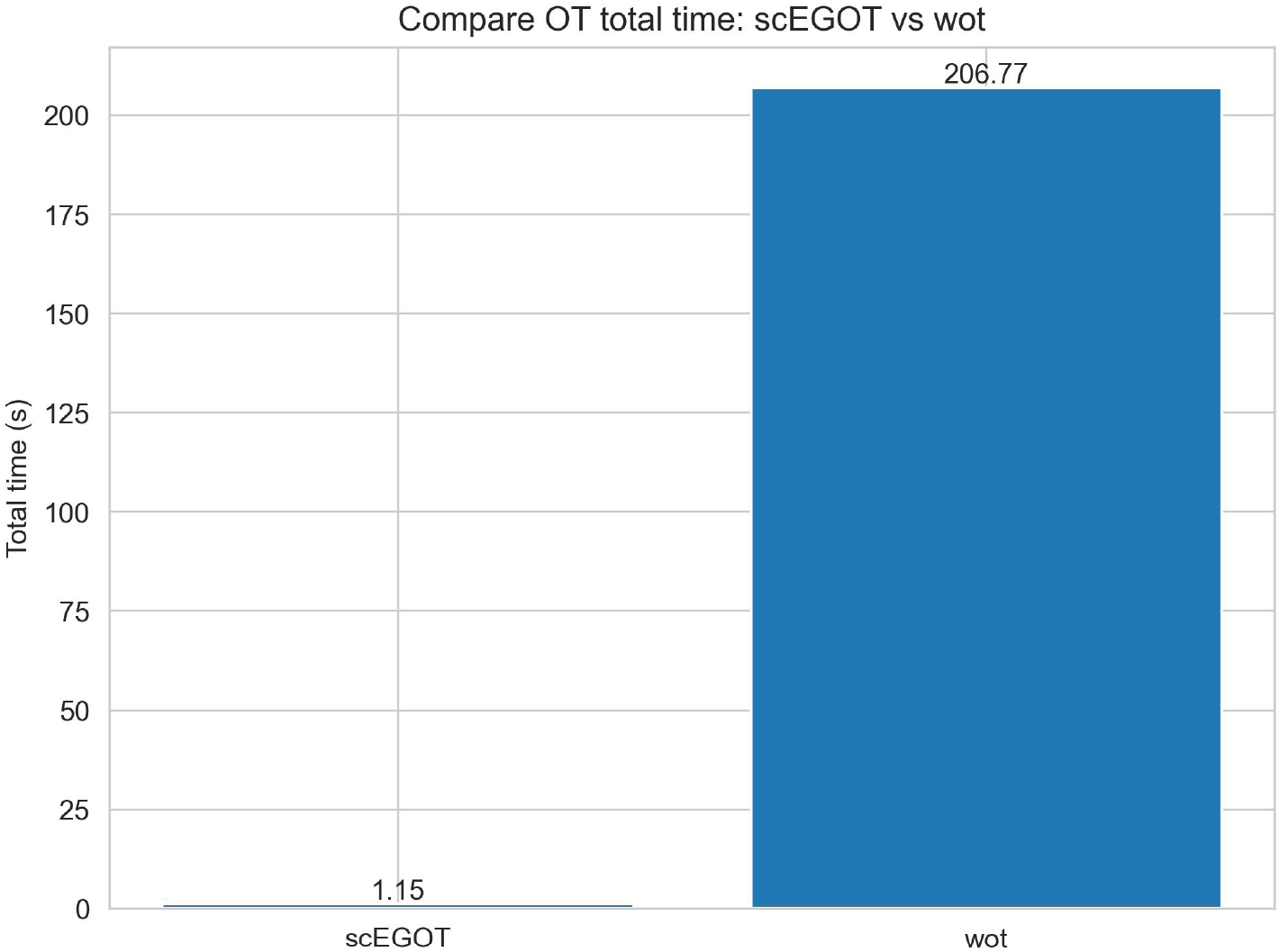
Comparison of the computational cost of optimal transport computation in scEGOT with that of Waddington-OT. When calculating the optimal transport for the entire time-series data of PGCLCs using the top 150 principal components. These experiments were performed on a MacBook Pro (14-inch, 2021) equipped with an Apple M1 Max and 64GB RAM.

**Figure S3:**
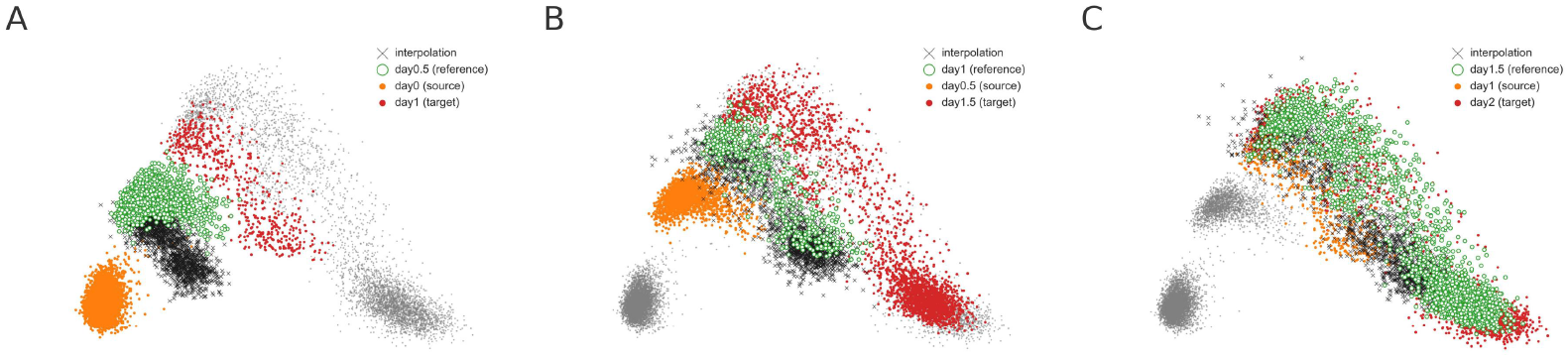
Comparison between the reference distribution and scEGOT interpolation distribution generated by datasets on all days on PCA coordinates. (A) Comparison between the reference distribution (day 0.5) and scEGOT interpolation distribution generated by datasets at days 0 and 1. (B) Comparison between the reference distribution (day 1) and scEGOT interpolation distribution generated by datasets at days 0.5 and 1.5. (C) Comparison between the reference distribution (day 1.5) and scEGOT interpolation distribution generated by datasets at days 1 and 2.

**Figure S4:**
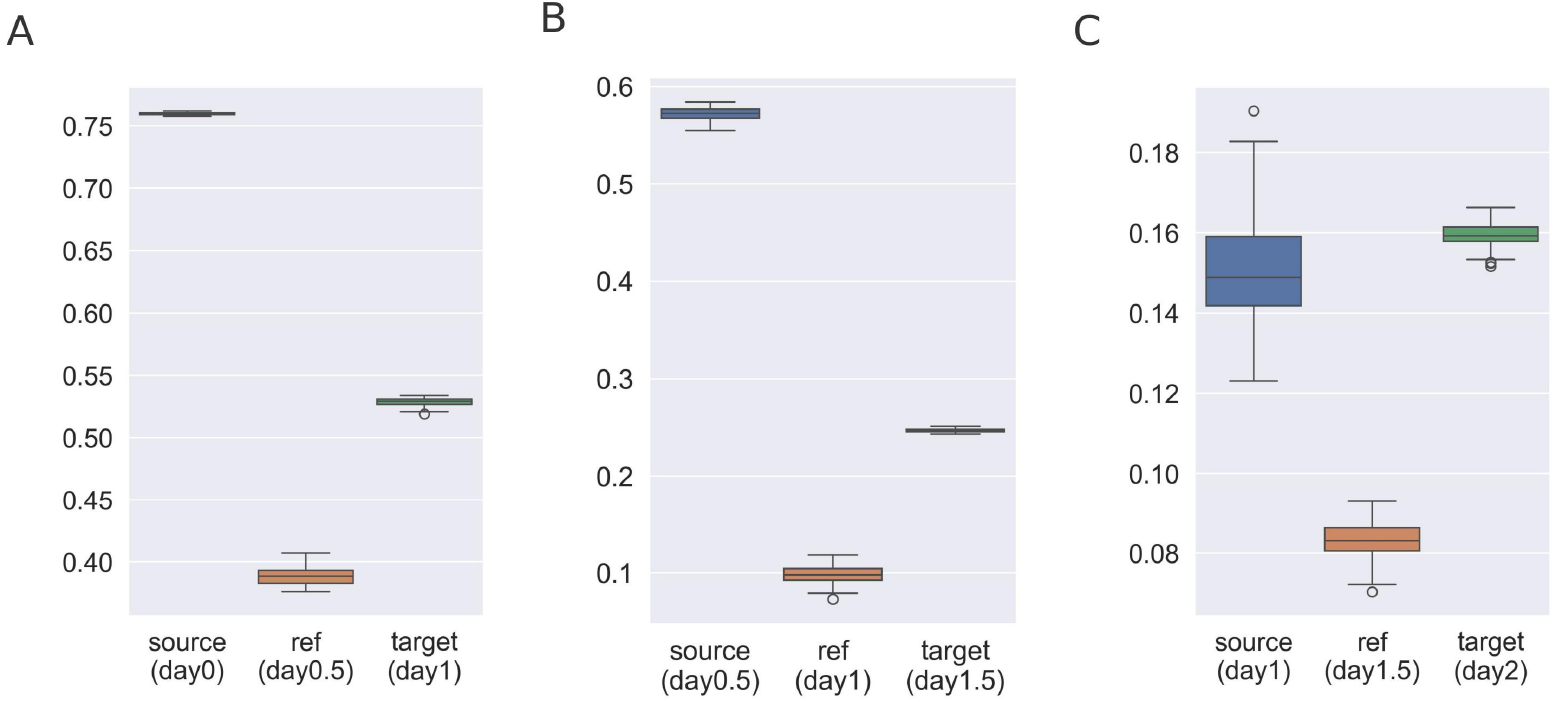
Box plot of the silhouette scores over 100 trials of the scEGOT interpolation versus source/reference/target; (A) day 0/day 0.5/day 1, (B) day 0.5/day 1/day 1.5, and (C) day 1/day 1.5/day 2.

**Figure S5:**
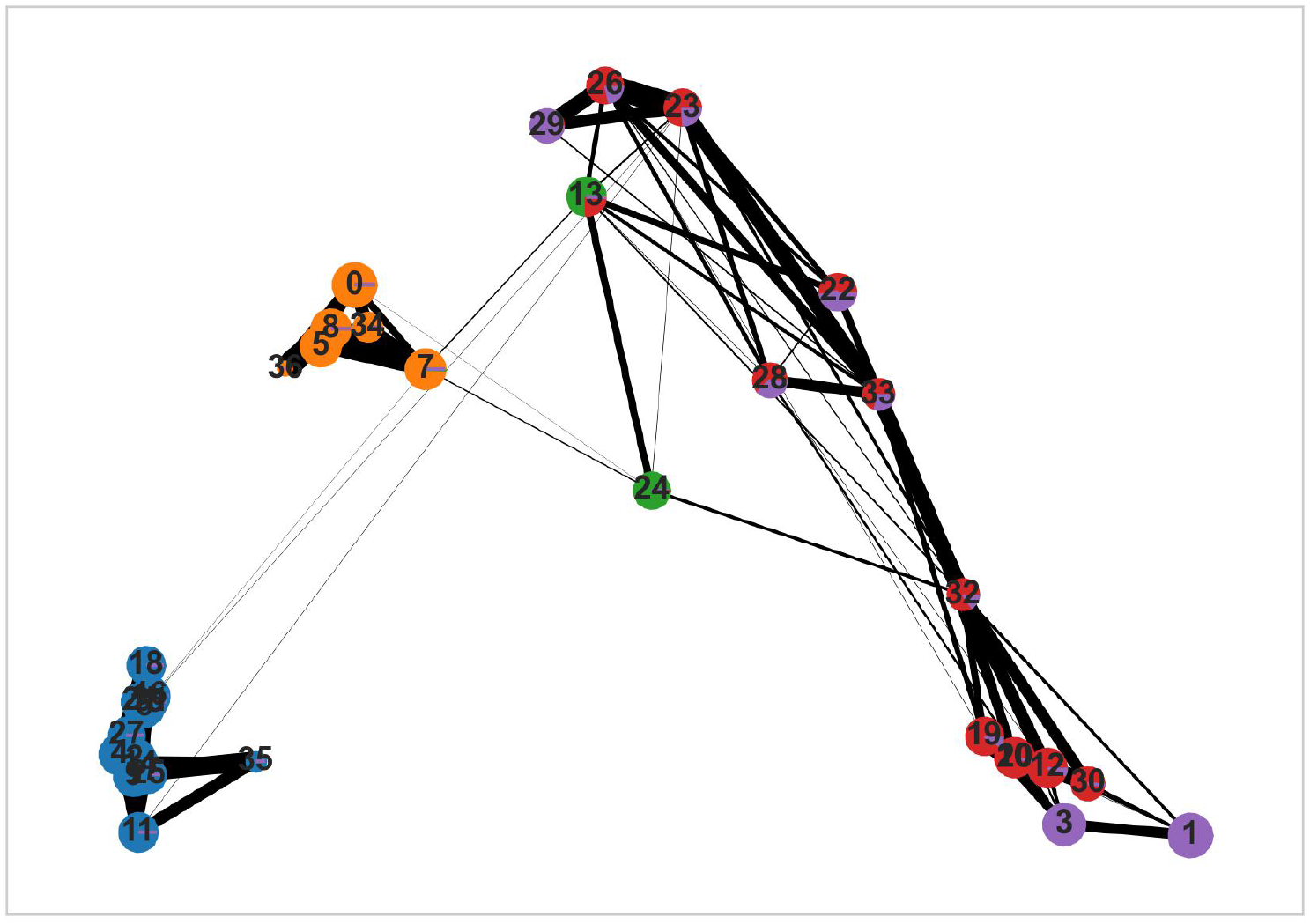
PAGA graph [4] applied to the human PGCLC induction dataset. The PAGA graph is represented as an undirected graph, showing the connectivity between different cell clusters.

**Figure S6:**
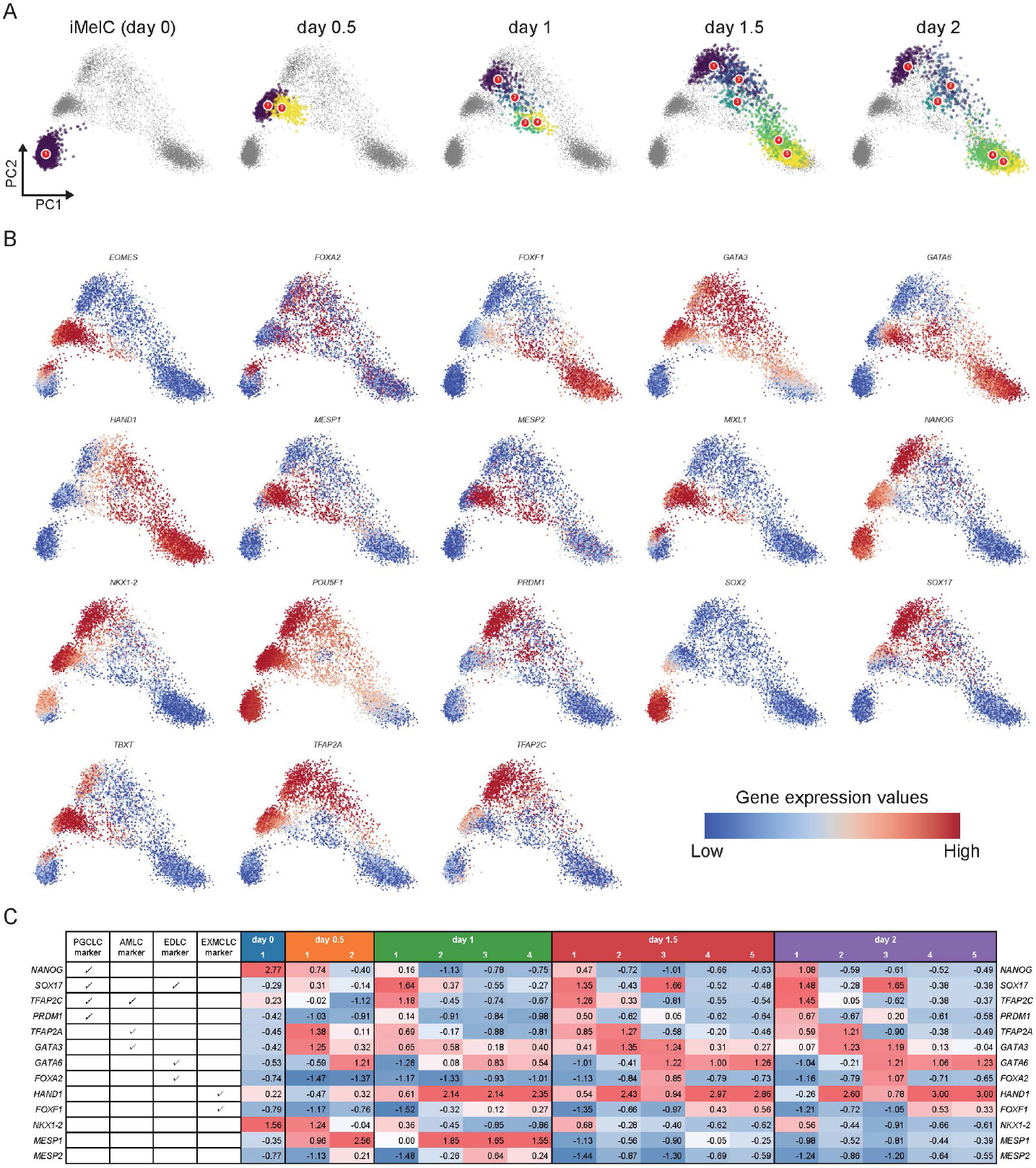
Clusters by the Gaussian mixture model (GMM) and marker gene’s expression values. (A) Clusters on PCA coordinate for each day. (B) Expression values of 18 marker genes. (C) Mean z-score of marker gene expression values for each cluster.

**Figure S7:**
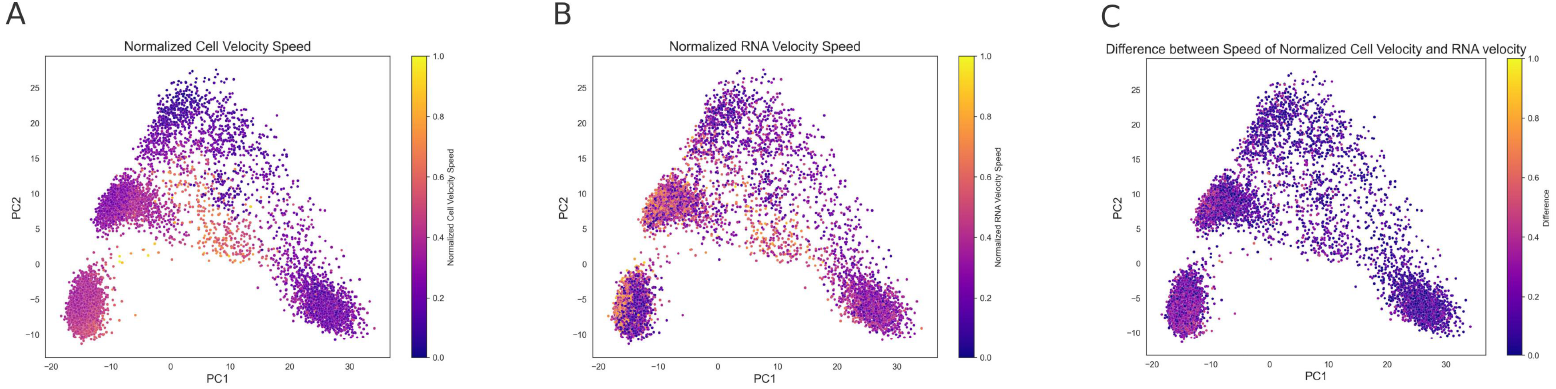
Comparison of normalized cell velocity and RNA velocity speeds. (A) Normalized cell velocity speed 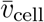, scaled to the range [0, 1]. (B) Normalized RNA velocity speed 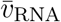, scaled to the range [0, 1]. (C) Difference between cell velocity and RNA velocity speeds, calculated as 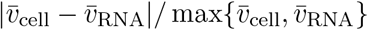

**Figure S8:**
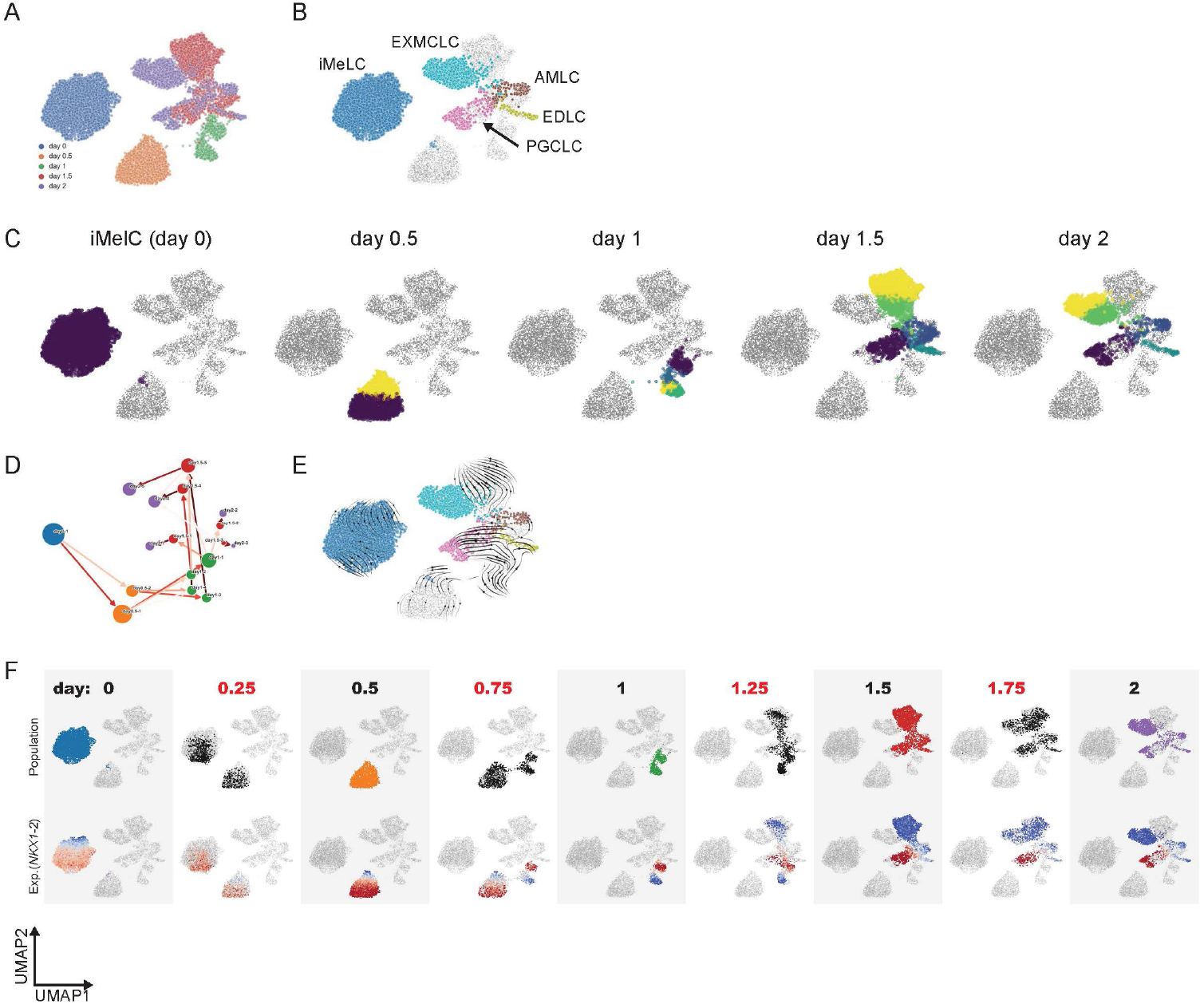
Visualization on the UMAP coordinate. UMAP plots are colored by (A) experimental days, (B) cell types, and (C) clusters by the Gaussian mixture model (GMM). (D), (E), (F) scEGOT’s outputs, cell state graph, cell velocity, and interpolation (top; cell population, bottom; *NKX1-2* expression value).

**Figure S9:**
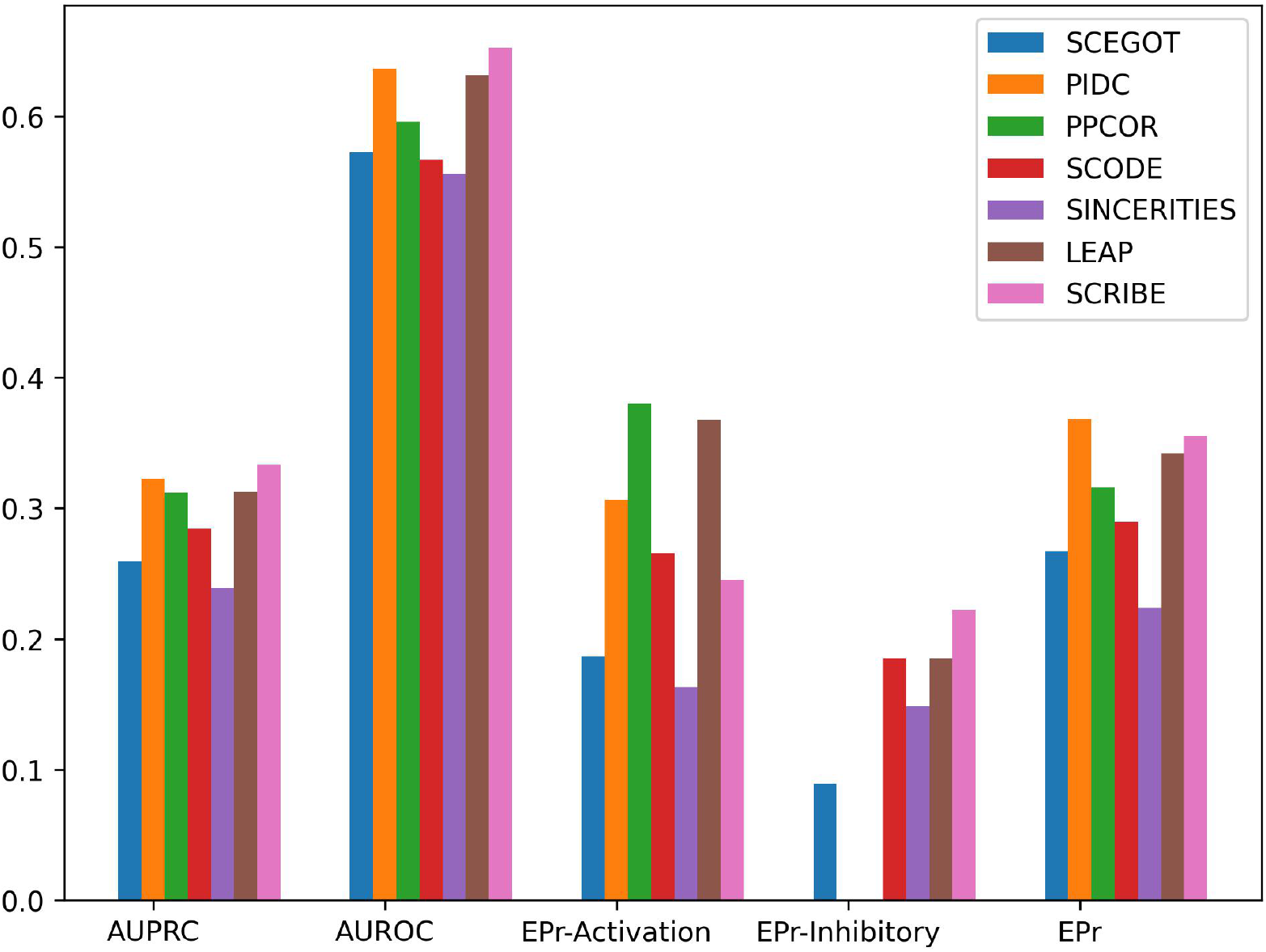
Comparison of scEGOT’s GRN inference algorithm to other methods for GRN inference using the BEELINE framework and its test data [5]. The performance was evaluated using the BLEval module from BEELINE. AUPRC (Area Under the Precision-Recall Curve): Measures the area under the precision-recall curve. AUROC (Area Under the Receiver Operating Characteristic Curve): Measures the area under the ROC curve. EPr (Early Precision): Calculates the precision within the top-k predicted edges, where k is the number of true edges. EPr-Activation: Calculates early precision specifically for activation-related edges. EPr-Inhibitory: Calculates early precision specifically for inhibition-related edges. When applying scEGOT, we first clustered the data based on pseudotime provided by the BEELINE benchmark test data and then treated these clusters as sequential time points.

**Figure S10:**
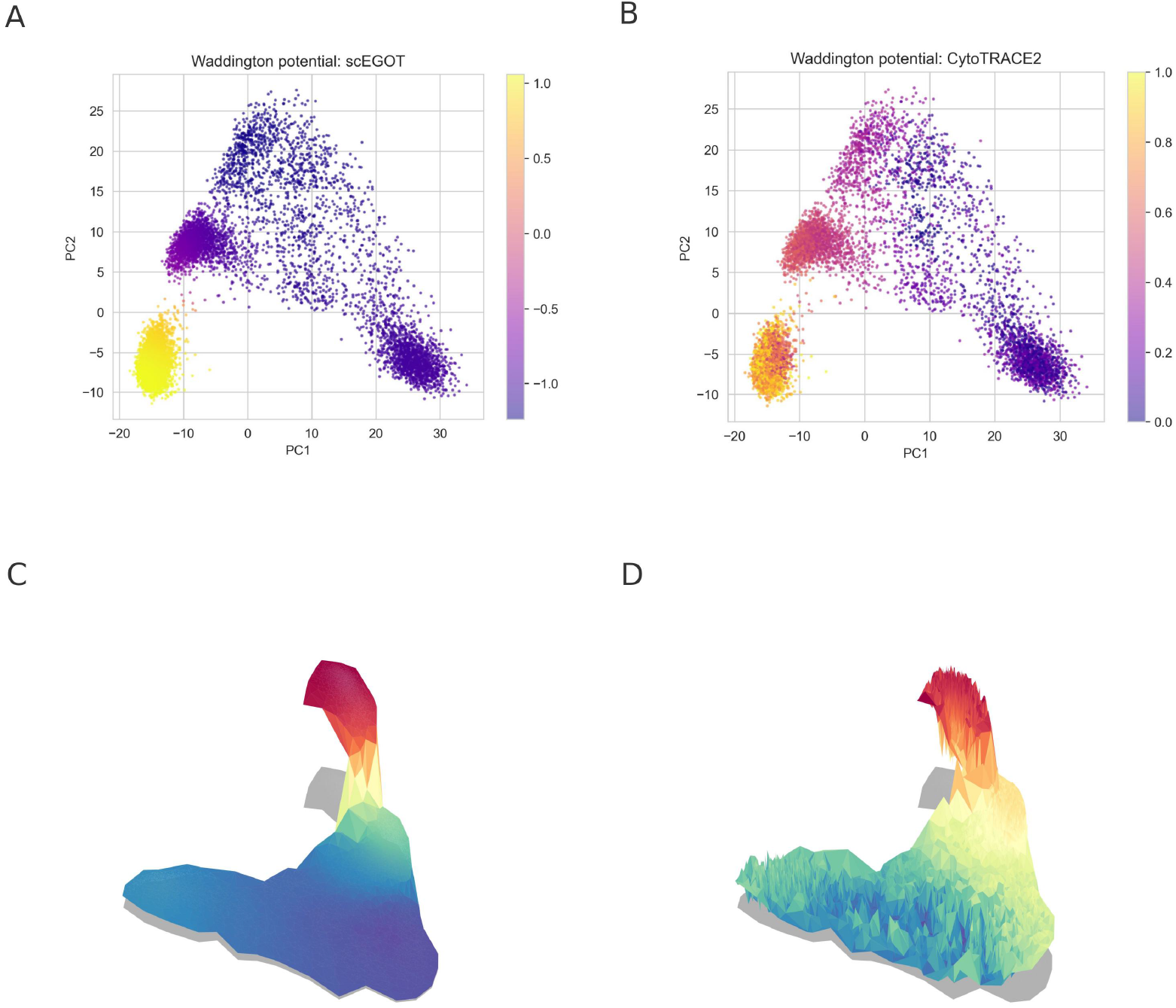
Comparison of the Waddington potential reconstructed by scEGOT ((A) 2-d, (C) 3-d) and the potential inferred by CytoTrace2 [6], which represents the potency score ((B) 2-d, (D) 3-d). We use (1−potency score) as the output of plots in CytoTrace2.

### Movies

Movie S1. Time-continuous gene expression dynamics of *NKX1-2* on the PCA coordinates. Movie S2. Time-continuous gene expression dynamics of *TFAP2A* on the PCA coordinates. Movie S3. Time-continuous gene expression dynamics of *NKX1-2* on the UMAP coordinates. Movie S4. Time-continuous gene expression dynamics of *TFAP2A* on the UMAP coordinates.

